# Dynamic signal processing by ribozyme-mediated RNA circuits to control gene expression

**DOI:** 10.1101/016915

**Authors:** Shensi Shen, Guillermo Rodrigo, Satya Prakash, Eszter Majer, Thomas E. Landrain, Boris Kirov, José-Antonio Daròs, Alfonso Jaramillo

## Abstract

Organisms have different circuitries that allow converting signal molecule levels to changes in gene expression. An important challenge in synthetic biology involves the *de novo* design of RNA modules enabling dynamic signal processing in live cells. This requires a scalable methodology for sensing, transmission, and actuation, which could be assembled into larger signaling networks. Here, we present a biochemical strategy to design RNA-mediated signal transduction cascades able to sense small molecules and small RNAs. We design switchable functional RNA domains by using strand-displacement techniques. We experimentally characterize the molecular mechanism underlying our synthetic RNA signaling cascades, show the ability to regulate gene expression with transduced RNA signals, and describe the signal processing response of our systems to periodic forcing in single live cells. The engineered systems integrate RNA-RNA interaction with available ribozyme and aptamer elements, providing new ways to engineer arbitrary complex gene circuits.

## Introduction

Natural signal transduction systems allow organisms to adapt to fluctuating environments, often by exploiting subcellular localization, molecular cascades and protein allostericity (1,2). A major challenge in synthetic biology involves the engineering of novel signaling systems that sense, process, and transmit information. Most engineering efforts have relied on the translational fusion of known protein domains with specific interaction or catalytic functionalities (2). However, this approach is limited by the availability of known natural interaction domains that are specific enough to avoid cross-talk with other molecules in the cellular context. Alternatively, the use of RNA as programmable molecules would allow engineering an unlimited number of interaction partners (3,4). This way, we propose to engineer synthetic signal transduction systems relying on RNA by using a transcriptional fusion strategy, exploiting sequence fragments with definite interaction and catalytic properties. In protein-based signaling, localized folding domains facilitate the engineering (or re-engineering) of multiple functions (5,6). Similarly, there are well-known RNA folding structures that are stable and capable to interact specifically with signaling molecules (aptamers) or to catalyze reactions (ribozymes) (4). In addition, the use of computational tools allows the prediction of conformational changes in many cases, opening the door to the engineering of signal transduction systems based on RNA (7). As a proof of concept, we here develop a system (to control gene expression with a molecular signal) that consists in the fusion of an aptazyme, acting as a molecular sensing element, with a riboregulator, acting as a signal mediator. To simplify the terminology, in the following we refer to this multifunctional RNA molecule as regazyme.

In this direction, pioneering work in synthetic biology inserted known aptamer domains into 5’ untranslated regions (UTRs) of messenger RNAs (mRNAs) to sense small molecules (10), and also exploited riboregulation in combination with small-molecule-responsive promoters to control gene networks and metabolic pathways (8,9). More recently, important steps towards RNA-based sensing have been carried out by engineering aptazymes in the 5’ or 3’ UTRs to sense both small molecules (11,12) and small RNAs (sRNAs) (13). Moreover, previous work has combined aptamers with riboregulators to create novel sensing devices (13-15). Those works exploit the programmability of RNA function through strand-displacement reactions and induced conformational changes. Here, our strategy allows engineering a one-to-two-component signal transduction system, where emerging RNA function is achieved by incorporating self-cleavage ability into a *trans*-acting riboregulator. This corresponds to a four-molecule system, where the first one is the signal molecule, either a small molecule or sRNA, and the last one is a *cis*-regulated mRNA as system’s readout. The other two molecules (two components) correspond to the sensor and mediator, which can be switched ON/OFF in presence/absence of the signal molecule, respectively.

The devised system shares properties with natural signaling systems (1). On the one hand, it is a one-component system from the input viewpoint. Thus, it has the advantage of subcellular localization independence. On the other hand, it is a two-component system from the output perspective (the sensor and mediator are different molecules after cleavage). Thanks to the modularity offered by the independence of the sensor and mediator domains, we could have a palette of domains with alternative functionalities. Compared to endogenous sensors (e.g., receptors), our sensor is not limited to the cell membrane, meanwhile the mediator (i.e., riboregulator) works like a phosphorylated transcription factor but at post-transcriptional level (interacting with a 5’ UTR rather than with a promoter). Our approach to engineer a signal transduction system combines the fusion of functional RNA elements together with the computational prediction of each conformational state. This is also possible with a protein-based system (16,17), although it could become much harder, requiring experimental screening towards appropriate transduction properties (5).

In this work, we show that RNA structure is predictable enough to allow a computational design strategy. In general, the tremendous size of the system’s sequence space prevents the *de novo* design without automation. We have previously demonstrated that an automated design methodology is able to generate *de novo* riboregulation in live cells (18). Therefore, we here propose to generalize such methodology to design RNA-mediated signal transduction systems. For that, we assume that any interaction between two RNAs is triggered by a seed (or toehold) sequence (18). In the case of a regazyme, the signal molecule induces a catalytic process that releases a riboregulator, which in turn induces a conformational change in the 5’ UTR that initiates interaction with the 16S ribosomal unit (18,19) in *Escherichia coli*. This way, we enforce a hierarchical mode of action consisting in switching ON each functional module, which is initially OFF.

In the following, we will provide a detailed description of the computational methodology to design a hierarchical system with functional RNA modules that couples molecular signals in the cell (either from the environment or from upstream biological systems) with gene expression. We will first describe the development of a methodology for nucleotide sequence design, and then we will present a mechanistic characterization to assess the self-cleavage activity of the regazyme. Subsequently, we will show results assessing signal transduction with time-dependent induction in bacterial cells, allowing the characterization of the dynamic regulatory properties at both population and single cell levels.

## Material and Methods

### Sequence design

We developed a Monte Carlo Simulated Annealing (20) optimization algorithm to design the transducer modules of regazymes provided the sequences of given aptazymes (or sRNA-induced ribozymes) and riboregulators (Supplementary Figure 6). For that, we constructed a basic energy model that involved three variables (to be minimized): the energy of activation corresponding to the catalytic activity of the aptazyme, the degree of accessibility of the riboregulator seed before cleavage, and the degree of obstruction of the seed after cleavage. The exposure or obstruction of the riboregulator seed is governed by secondary structure, but the aptazyme involves tertiary contacts. We here simplified the problem by only considering the secondary structure of the aptamer to calculate the energy of activation for cleavage. Rounds of random mutations (replacements, additions or deletions) were applied and selected with the energy-based objective function. We used the Vienna RNA package (21) for energy and structure calculation (see further details in **Supplementary Materials and Methods**). The sequences of the engineered regazymes in this work are shown in Supplementary Tables 1 - 3.

### Plasmid construction

The different RNA devices were chemically synthesized and cloned in plasmid pSynth (pUC replication origin, ampicillin resistance marker) and then subcloned into plasmids pSTC1 or pSTC2. These two plasmids contain a pSC101m replication origin (a mutated pSC101 ori giving a high copy number) and a kanamycin resistance marker (Supplementary Figures 1, 2). The pSTC2 vector is based on our previously reported vector pSTC1 (18) by removing the mRFP coding sequence and tagging the carboxyl terminus of the superfolder GFP (sfGFP) (22) with the *ssr*A degradation tag (23). Dysfunctional regazymes were constructed by PCR-based site-directed mutagenesis (see **Supplementary Materials and Methods**). Strains and plasmids used in this study are listed in Supplementary Table 6.

### Intracellular catalytic activity

Processing extent of regazyme at different time points (0, 2, 4, 8, 16 and 32 min) was analyzed by northern blot hybridization using a complementary [^32^P]-labelled RNA probe after separating the different RNA samples by denaturing polyacrylamide gel electrophoresis (PAGE). RNA preparations were mixed with formamide loading buffer for denaturation, followed by PAGE separation in 5% polyacrylamide gels including 8 M urea and TBE buffer. Gels were stained with ethidium bromide. Membranes were hybridized overnight, imaged by autoradiography, and then hybridization signals quantified by phosphorimetry (Fujifilm FLA-5100). See more details in **Supplementary Materials and Methods**.

### Fluorescence quantification

Cells were grown overnight in LB medium, and were refreshed in culture tubes with LB medium in order to reach stationary phase. Cells were then diluted 1:200 in 200 μL of M9 minimal medium in each well of the plate (Custom Corning Costar). The plate was incubated in an Infinite F500 multi-well fluorometer (TECAN) at 37 °C with shaking. It was assayed with an automatic repeating protocol of absorbance measurements (600 nm absorbance filter) and fluorescence measurements (480/20 nm excitation filter - 530/25 nm emission filter for sfGFP) every 15 min. All samples were present in triplicate on the plate (see further details in **Supplementary Materials and Methods**).

### Single cell microfluidics analysis

The design of our microfluidics device (Supplementary Figure 16) (24), which was performed in AUTOCAD (AUTODESK), was adapted from the previous one reported by Hasty and coworkers (25). All images were acquired using Zeiss Axio Observer Z1 microscopy (Zeiss). The microscope resolution was 0.24 μm with Optovariation 1.6X, resulting total magnification 1600X for both bright field and fluorescent images. Images were analyzed with MATLAB (MathWorks). Cells were tracked by defining a cell-to-cell distance matrix and the cell lineages were reconstructed. Finally, the fluorescence level of each cell in each fluorescence frame was extracted (see further details in **Supplementary Materials and Methods**).

## Results

### Computational design of RNA-mediated signal transduction systems to control gene expression

Our modular strategy consists in designing switchable functional RNA domains, which is implemented by exploiting strand-displacement principles together with the engineering of allosteric conformational states. This way, we can design chains of several domains that are activated in cascade. Without loss of generality, we considered a system composed of two transcriptional units: regazyme and mRNA of a reporter gene (e.g., a gene coding for a green fluorescent protein –GFP), but our methodology could be generalized to an arbitrary number of transcriptional units containing switchable functional elements. To engineer such a synthetic RNA system implementing the transduction of molecular signals into changes in gene expression, we took advantage of a standard physicochemical model (based on Watson-Crick and wobble pairing) predicting RNA secondary structure and free energy (26) to be used in an optimization algorithm to select for the hierarchical activation of functional RNA modules in the cascade (7).

In particular, our system corresponds to a cascade of three modules: sensor (aptazyme designed to specifically respond to a given ligand), mediator (riboregulator designed to specifically activate a *cis*-repressed ribosome-binding site –RBS), and actuator (mRNA with *cis*-repressed RBS) (Figure 1a). To create the regazyme, we fused an aptazyme element, acting as a molecular sensing device, with a riboregulator, acting as a signal mediator, into the same transcriptional unit. This fusion is performed with flanking sequences that form a stem and function as a transducer module, in the same way as when designing allosteric aptamers (27). The sensor domain (aptazyme) is initially in a state OFF (catalytically inactive) and is switched to ON (catalytically active) only when it acquires its functional conformation, which is induced by the signal molecule. The mediator domain (riboregulator) will be in a state ON when its seed sequence is exposed to the solvent. We designed the transducer module to ensure that the sensor and mediator were ON/OFF in presence/absence of the signal molecule. This prevents any premature release of the mediator or any direct activation of gene expression by the regazyme. Afterwards, the input signal produces a stabilization of an alternative conformation where the aptazyme is active. Once the aptazyme is active, it will self-cleave releasing the mediator domain, which is then switched on (i.e., the riboregulator seed sequence becomes exposed). Once the mediator domain is active, it will diffuse towards its target genes (in particular, to interact with 5’ UTRs), similarly to phosphorylated transcription factors in the conventional two-component systems (1). To be noted, the independence between the sensor and mediator domains favors expanding the functional repertoire, which allows them to be exchanged with alternative domains.

**Figure 1.**
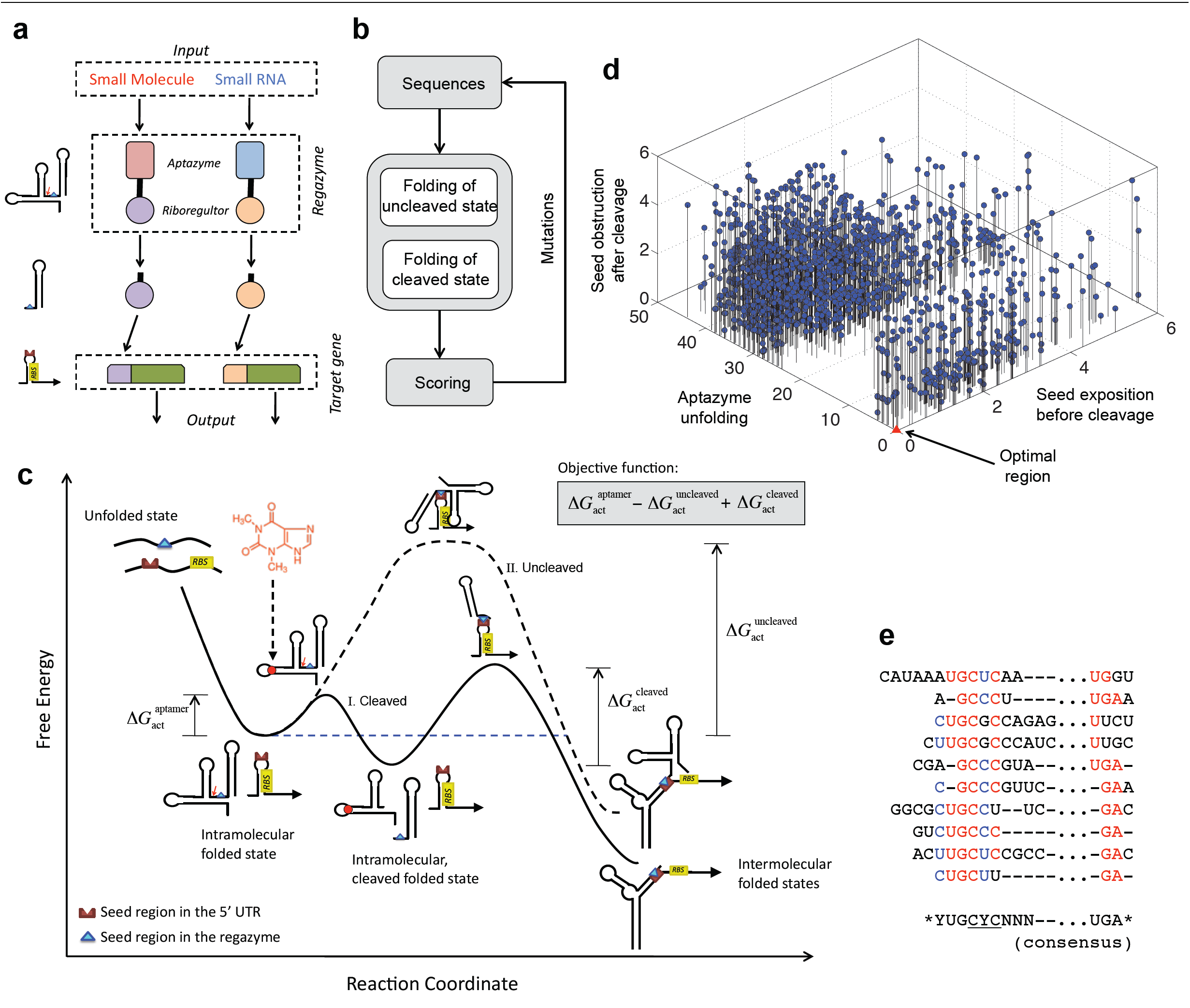
Computational design of the regazyme signaling pathway. (a) Scheme of the modular system, where a signal molecule (either a small molecule or a sRNA) induces a catalytic reaction that releases a riboregulator able to activate gene expression. Each signaling pathway is viewed as a wire carrying information. (b) Scheme of the optimization loop, where a regazyme sequence is iteratively mutated and evaluated according to an objective function. (c) Energy landscape of the signaling pathway showing the different conformational states (intra- and intermolecular), together with the three free energy terms of the objective function, in terms of a reaction coordinate. Solid line illustrates the trajectory corresponding to the ligand-induced cleavage of the regazyme and subsequent binding of the riboregulator to the mRNA. Dashed line corresponds to the trajectory where the uncleaved regazyme binds to the mRNA. (d) Computation of the free energy terms of the objective function for 1,000 random sequences to evaluate their distribution. (e) Alignment of different optimized sequences with aptazyme theoHHAz and riboregulator RAJ12. Highly conserved nucleotides are highlighted in red or blue, and the consensus sequence is shown.

We here propose a new methodology to engineer one-to-two-component signal transduction, which combines the advantages of subcellular independence of one-component systems and of modularity of two-component systems. We constructed a combinatorial optimization problem to explore the sequence space of the transducer module (Figure 1b), where a nucleotide-level energy model considering the conformational states (uncleaved and cleaved) of the regazyme was used to evaluate the performance of the generated sequences. For each state, the model accounts for its free energy and its secondary structure. As objectives to be optimized (computed as Hamming distances), the algorithm considers the energy of activation corresponding to the catalytic activity of the aptazyme (which we assume depends on the correct formation of the aptamer in the uncleaved state), and the degree of exposure to the solvent of the riboregulator seed before and after cleavage (28). Figure 1c illustrates the energy landscape associated to the molecular mechanism of the regazyme, reporting the different conformational states and their corresponding free energy levels (see also Supplementary Figure 4). The reaction coordinate was defined here as the number of intermolecular hydrogen bonds, on one side, between the ligand and the aptazyme and, on the other side, between the riboregulator and the 5’ UTR (in terms of base-pairs). In absence of signal molecule, the progression of the reaction is limited by the presence of a high-energy intermediate that prevents the interaction between the regazyme and the 5’ UTR. However, when the signal molecule is at sufficient concentration, a cleavage is produced and then the activation energy for the resulting riboregulatory element is lowered, which speeds up the reaction (29).

As shown by a random sampling of 1,000 sequences (Figure 1d), an optimal score (zero, as our score is considered as a penalty) is very unlikely to be obtained arbitrarily. This means that this is a difficult design problem for a manual approach, requiring automated computation for efficient sequence design. Our algorithm designs by optimization the sequences implementing the intended signal transduction according to the objective function. Even though distinct solutions can be equally good computationally (i.e., according to the objective function), experiments could distill differences in performance among them. We observed, for the sampled sequences, that the aptamer (in the uncleaved state) is correctly formed only for a small subset of sequences (Supplementary Figure 5). Moreover, the resulting distribution is apparently bimodal (Sarle’s bimodality coefficient *BC* = 0.630 > 5/9) (30), which may be explained by an all-or-none formation of the functional structure of the aptazyme. Such pre-organized conformations will favor ligand binding and subsequent cleavage, whereas structures requiring considerable rearrangements will be offside due to a given free energy barrier (29). The distribution of score values along the axis representing the seed exposure in the uncleaved state is more homogeneous (*BC* = 0.467 < 5/9), whereas the distribution in the cleaved state shows substantial heterogeneity (*BC* = 0.639 > 5/9). This may be explained by an interaction of the seed region with part of the 5’ end after cleavage (see for example Supplementary Figure 8).

### Modularity in the design of regazymes

In this work, we considered three possible sensor domains, two sensing a small molecule (theophylline –Theo– and thiamine pyrophosphate –TPP, Figure 2 and Supplementary Figure 7), and another sensing a specific sRNA (Break1, Figure 3). This sRNA is induced with anhydrotetracycline (aTc) in our system. More specifically, each sensor is composed of a binding domain (e.g., an aptamer) and a catalytic domain (e.g., a hammerhead ribozyme). Our ligand-induced ribozymes (aptazymes) are theoHHAz and tppHHAz for sensing small molecules (11,31), and breakHHRz for sensing sRNA (32) (Supplementary Figure 3). For the mediator domain, we considered three synthetic riboregulators known to activate the initiation of translation, two engineered in Rodrigo *et al.* (RAJ11 and RAJ12) (18) and one in Isaacs *et al.* (RR12) (19) (Supplementary Figure 8). We then designed the rest of the regazyme sequence according to the specifications required to generate the RNA signaling cascade. Exploiting the modularity of this system, we engineered the following regazymes: theoHHAzRAJ11, theoHHAzRAJ12, theoHHAzRR12, tppHHAzRAJ12, and breakHHAzRAJ12. The regazyme produces a mediator molecule (riboregulator) that is independent of the signal and sensor molecules. In the following, we investigate, on the one hand, how different signal molecules (Theo, TPP and Break1) activate a common mediator (RAJ12), and, on the other hand, how different implementations of the wire (RAJ11, RAJ12 and RR12) transduce the information from a common signal molecule (Theo).

**Figure 2.**
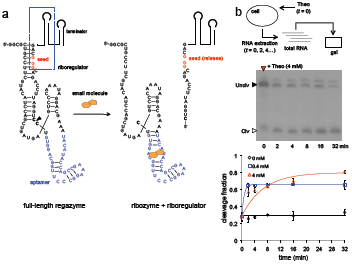
Molecular characterization of small-molecule-sensing regazyme. (a) Sequence and structure of the regazyme theoHHAzRAJ12. A small molecule (Theo) binds to the regazyme to reconstitute the active conformation of the ribozyme and then produce the cleavage. An arrow marks the cleavage site, between the transducer module and the ribozyme core. The seed of the riboregulator is paired in the suncleaved state. (b) Time-dependent electrophoretic analysis of cellular RNA extracts taken at different time points; gel shown for 4 mM Theo. Quantification of dynamic RNA processing for different concentrations of the signal molecule (Theo). Data fitted with a generalized exponential decay model with production, where the temporal factor is (1-exp(-*λt*))^m^, with *m* ≈ 1. Error bars represent standard deviations over replicates.

**Figure 3.**
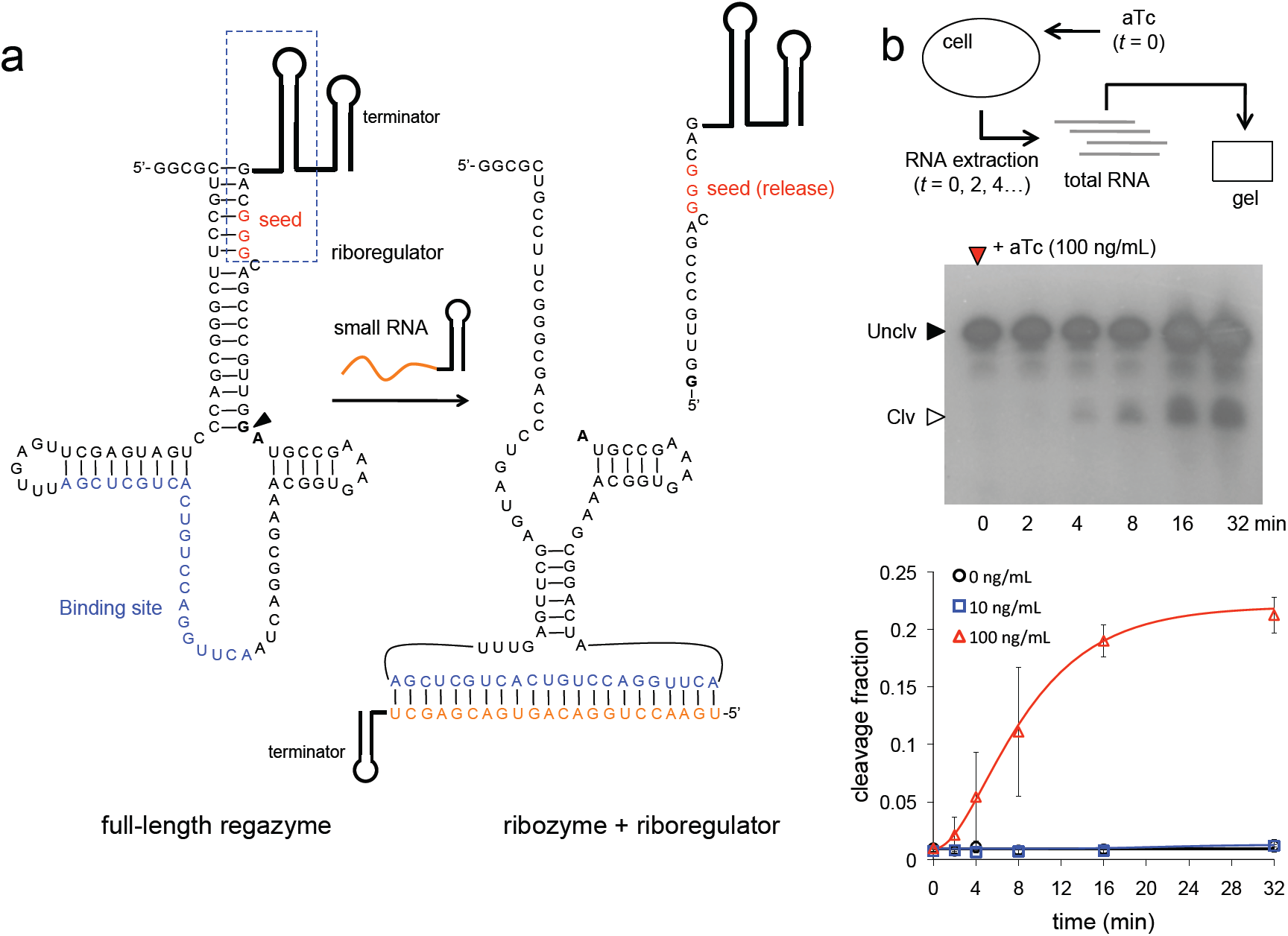
Molecular characterization of sRNA-sensing regazyme. (a) Sequence and structure of the regazyme breakHHRzRAJ12. A sRNA binds to the regazyme to reconstitute the active conformation of the ribozyme and then produce the cleavage. An arrow marks the cleavage site, between the transducer module and the ribozyme core. The seed of the riboregulator is paired in the uncleaved state. (b) Time-dependent electrophoretic analysis of cellular RNA extracts taken at different time points; gel shown for 100 ng/mL aTc. Quantification of dynamic RNA processing for different concentrations of the signal molecule (aTc). Data fitted with a generalized exponential decay model with production, where the temporal factor is (1-exp(-*λt*))^m^, with *m* ≈ 2. Error bars represent standard deviations over replicates.

Our computational approach allowed us to investigate the designability (defined as the number of sequences that have the desired biochemical function) of the solution space for a particular couple of aptazyme and riboregulator. We expect, nevertheless, a higher designability when sequences are allowed to vary in length, as it is the case for our algorithm. It is instructive to align multiple solutions to reveal conserved nucleotide positions. Figure 1e shows, for different designs, the consensus sequence for a given choice (aptazyme theoHHAz and riboregulator RAJ12). Sequences (corresponding to the 5’ and 3’ regions of the aptazyme, see Supplementary Figure 6a) were aligned by using the anti-seed consensus sequence (CYC in this case; note that the seed sequence is GGG) as reference. In addition, this modularity would allow a hierarchical design of the regazyme molecule. To create a suitable pipeline, we can exploit computational algorithms to: *i*) design the binding domain (aptamer in particular) for a specific signal molecule (33), *ii*) design the riboregulator and cognate 5’ UTR (18), and *iii*) apply the methodology developed in this work to design the appropriate transducer module. Experimental screenings or directed evolution techniques (34) could also be applied, especially to link the binding and catalytic domains (see discussion).

### Molecular characterization of RNA-mediated signal transduction

To analyze the mechanism of the signaling pathway, we first carried out a kinetic and dose-dependent study of the catalytic activity. The predicted secondary structure of the small-molecule-sensing regazyme theoHHAzRAJ12 in the uncleaved state (Figure 2a) shows, as designed, that the aptamer is already arranged for Theo sensing, and that the seed region of the riboregulator is blocked by the transducer module. After cleavage at the CC dinucleotide site between the transducer module and the ribozyme core (Figure 2a, marked by an arrow), the seed region is released, which allows the riboregulator to interact downstream with the cognate 5’ UTR of the reporter gene. An analogous seed-based structural mechanism is illustrated for the sRNA-sensing regazyme breakHHRzRAJ12 (Figure 3a). In this case, the binding domain is only partially paired to allow an efficient interaction with the signal sRNA (Break1), and the cleavage is done at the GA dinucleotide site (Figure 3a, marked by an arrow). Indeed, there is a seed-mediated interaction between Break1 and the regazyme, similar to the interaction between the riboregulator and the 5’ UTR. Of note, our regazyme breakHHRzRAJ12 implements for the first time an RNA cascade in live cells (independent of any protein-based machinery).

To monitor the dynamic RNA processing of the system, we performed a gel assay from cellular RNA extracts. Cells expressing regazyme theoHHAzRAJ12 or breakHHRzRAJ12 were induced with different concentrations of Theo or aTc and lysed at several time points. The gel assays in both cases showed fast dynamic RNA processing, reaching steady states in almost 16 min. The observed cleavage rate (fitted with a model of exponential decay with production) is 0.15 min^−1^ for theoHHAzRAJ12 with 4 mM Theo, although the model does not capture finely the experimental trend (Figure 2b **and** Supplementary Figure 9a). In this case, the band corresponding to the released riboregulator (of 114 nt, accounting for the terminator) migrates faster than expected. This band was not observed without Theo, indicating that indeed it is a product of the cleavage reaction. Moreover, we note that our probe did not detect the 5’ fragment after cleavage, suggesting a fast degradation of this new species. In those conditions, the maximal cleavage fraction is about 80%, which is more than 2.5-fold increase with respect to the basal state (almost 30%). With 0.4 mM Theo, the observed cleavage rate is 1.5 min^−1^ with a more accurate fitting (note that the discrepancy between the rates at 4 and 0.4 mM is indeed due to the model fitting; at 4 min the fraction cleaved is about 60% in both cases). According to previous experimental results *in vitro* without RNA production and degradation (11), the observed cleavage rates of theoHHAz in absence and presence (4 mM) of Theo are 1.3 min^−1^ and 3.6 min^−1^, respectively, with a maximal cleavage fraction of 90%. Certainly, the dynamic response *in vivo* faces additional challenges due to the balance between production and degradation. For breakHHRzRAJ12, the observed cleavage rate is 0.17 min^−1^ with 100 ng/mL aTc, but no activity is reported for lower concentrations of this inducer (Figure 3b **and** Supplementary Figure 9b). Here, the band corresponding to the released riboregulator (of 112 nt, also accounting for the terminator) migrates slower than expected, although it was not observed (as before) without aTc, indicating that indeed it is a product of the cleavage reaction. These anomalous migrations could be due to a difference in expected length (e.g., unpredicted transcription termination, as most of the cleavage occurs co-transcriptionally) or to a residual structure in the released riboregulator even after using 8 M urea in the gel, among other possibilities. In case of breakHHRzRAJ12, previous assays *in vitro* (32) revealed a rate of 0.11 min^−1^ with single-stranded DNA as ligand (3 μM). The maximal cleavage fraction reported here is almost 25% (about 1% for the basal state), which shows a big discrepancy with those previous *in vitro* results (about 75%). One possible explanation is that the expression of the sRNA with 100 ng/mL aTc does not saturate the system, because the regazyme is expressed from a strong constitutive promoter, and also because of a high effective dissociation constant. Of relevance, this regazyme has much lower leakage (1% versus 30%, although maintaining similar fold-changes), which could be important in case of sensitive systems. Moreover, we observed higher heterogeneity in the dynamic response (from cell to cell) for this sRNA-sensing regazyme, which could be a result of a heterogeneous expression of the sRNA or even of a certain heterogeneous sRNA-regazyme interaction. As a result, by predicting RNA structures and quantifying cellular RNA extracts, we have shown the precise signal sensing and subsequent cleavage to release a functional riboregulator.

To further confirm that our devices behave as expected, we performed *in vitro* transcription of systems theoHHAzRAJ12 and breakHHRzRAJ12. The experiments showed similar cleavage fractions (with respect to the *in vivo* assays) after 30 min of reaction (Supplementary Figure 10a). For theoHHAzRAJ12, 70% of the molecules were cleaved *in vitro*, whereas 80% were *in vivo*. For breakHHRzRAJ12, 25% of the molecules were cleaved both *in vitro* and *in vivo*. We also observed that theoHHAzRAJ12 was cleaved in higher extent in absence of ligand. Because *in vitro* we can neglect degradation, the cleavage fraction is expected to increase with time, in presence of ligand and also in absence of it due to the basal activity of the ribozyme. We further performed a time-course assay to study the cleavage of the regazymes. As shown in our experimental results, the fraction of cleaved products of theoHHAzRAJ12 in test tubes was only about 10% larger when Theo was present (Supplementary Figure 10b). However, when the same RNA was monitored in bacterial cells, the apparent level of induction of cleavage activity by theophylline was more than 2.5-fold. The difference in cleavage *in vitro* in case of breakHHRzRAJ12 with respect to the presence or not of Break1 (introduced as DNA oligo) was more remarkable (Supplementary Figure 10c). There is certain number of reasons why an RNA might exhibit different behaviors *in vitro* than *in vivo*. For example, *in vitro* the regazyme might exist in thermodynamic equilibrium with its ligand, resulting in a different effective dissociation rate, or differ slightly in length from the strands *in vivo*. In addition, *in vitro* we used T3 polymerase for transcription (without terminators) instead of *E. coli* polymerase and a higher Mg^2+^ concentration than *in vivo*, which might result in differences in folding and cleavage kinetics of the regazyme. In the following, we present the net effect of regazyme cleavage and riboregulator release on GFP expression *in vivo* (both at the population and single cell levels) with and without the ligand, showing a regulatory behavior as designed.

### Regulation of gene expression in live cells with transduced RNA signal

To characterize the dynamic range of our engineered systems, we placed the transcriptional units corresponding to the regazyme and mRNA of the GFP reporter gene under the control of tunable promoters (35). These promoters can be induced with isopropyl-β-D-thiogalactopyranoside (IPTG) and aTc in *E. coli* cells expressing constitutively the repressors LacI and TetR. Thus, our systems implement multi-input AND logic circuits (Figures 4a). Moreover, a control system was implemented by using a dysfunctional mutated regazyme (Figure 4b). In the implementation for small-molecule signaling, aTc and IPTG control the expression of the regazyme and the mRNA, and Theo is the signal molecule that induces the cleavage of the regazyme to release the riboregulator. Figure 4c shows the fluorescence results of the system based on regazyme theoHHAzRAJ12 for all possible combinations of inducers (IPTG, aTc and Theo). The observed weak activation of fluorescence in absence of the signal molecule, but in presense of IPTG and aTc, can be explained by the leakage of self-cleavage of the regazyme (see also Supplementary Figure 11). The dysfunctional regazyme, obtained by a two-nucleotide mutation in the aptamer domain that abolishes ligand binding (Supplementary Figure 7a), was shown to significantly decrease GFP expression (Figure 4c). Furthermore, a single-nucleotide mutation in the ribozyme catalytic core (theoHHAzRAJ12AGm, A to G in Supplementary Figure 7a) (46) that inhibits the self-cleavage activity showed decreased GFP expression (Supplementary Figure 11c). An additional inactivating point mutation (theoHHAzRAJ12Cm and theoHHAzRR12Cm, U to G in Supplementary Figure 7a) (11) also revealed decreased GFP expression with respect to the native sequence (Supplementary Figures 11a,b). We also engineered and characterized systems based on riboregulators RR12 (Figure 4d) and RAJ11 (Supplementary Figure 11d), although the riboregulatory activity of theoHHAzRAJ11 with respect to its dysfunctional mutant was more moderate. Furthermore, these three regazymes for Theo-signaling have responsiveness in a dose-dependent manner (Supplementary Figure 14) with an effective dissociation constant of about 1 mM. Another engineered system for TPP signaling (regazyme tppHHAzRAJ12) showed no significant riboregulatory activity (Supplementary Figure 15).

**Figure 4.**
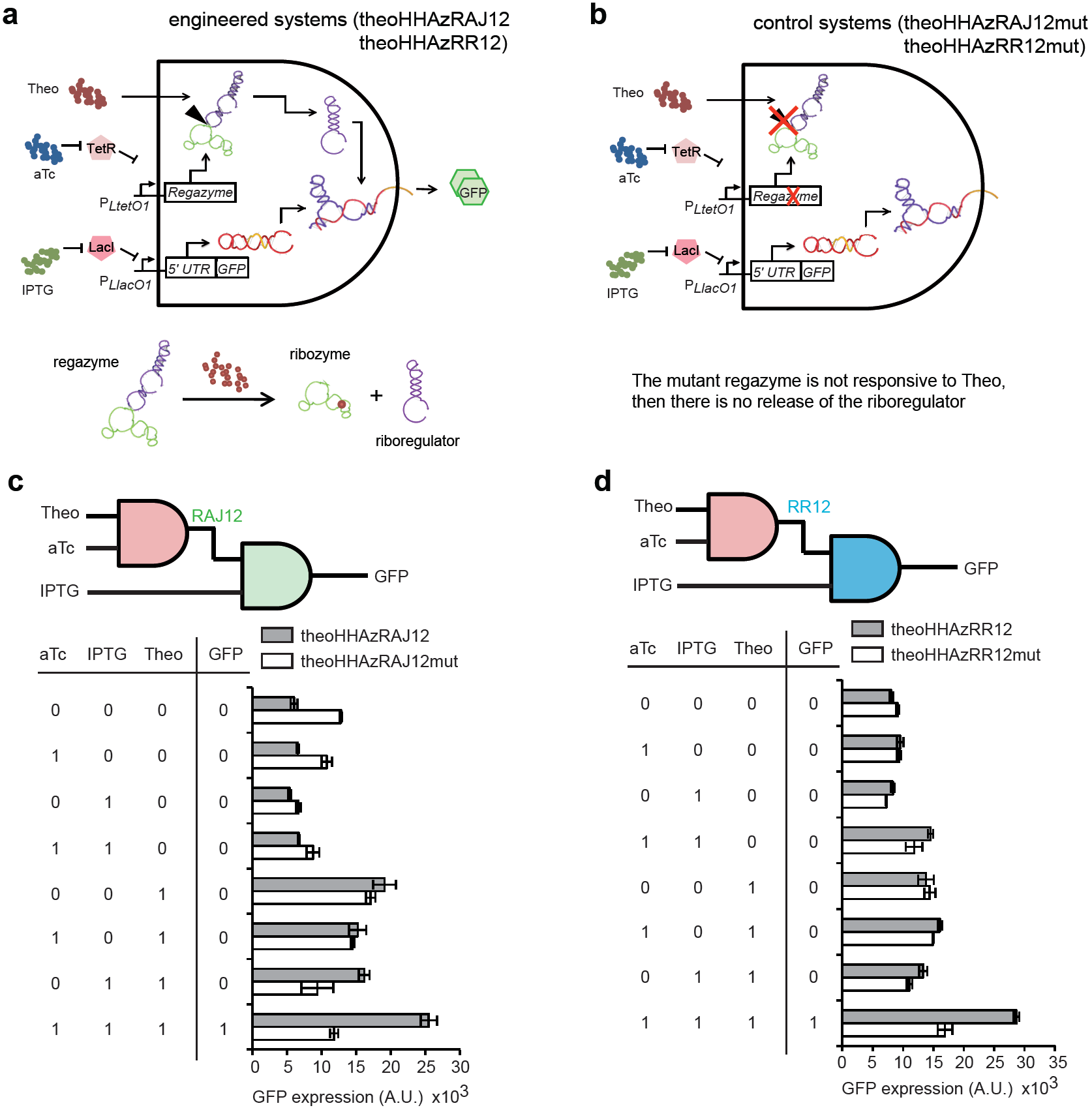
Functional characterization of small-molecule-sensing regazymes. (a, b) Schemes of the engineered RNA-based circuit to sense a small molecule and its corresponding control. (c, d) Digital scheme, associated Truth table, and fluorescence results of regazymes theoHHAzRAJ12 and theoHHAzRR12 (gray bars), and of their dysfunctional mutants (white bars) for all possible combinations of inducers. Error bars represent standard deviations over replicates.

In the implementation for sRNA signaling, aTc and IPTG control the expression of the sRNA working as signal molecule and the mRNA, whereas the regazyme is expressed from a strong constitutive promoter (Figure 5a). In this case, the cleavage of the regazyme is induced by that sRNA. Figure 5c shows the fluorescence results of the system based on regazyme breakHHRzRAJ12 for all possible combinations of inducers (IPTG and aTc). This logic circuit could further be expanded to integrate more inputs by replacing the constitutive promoter of the regazyme to other tunable promoter. To exclude the possibility that the sRNA Break1 could directly activate the *cis*-repressed reporter gene, we generated a control system (breakRAJ12) by using a dysfunctional mutant removing the regazyme element (Figure 5b and Supplementary Figure 13), revealing no significant riboregulatory activity in this case (Figure 5c).

**Figure 5.**
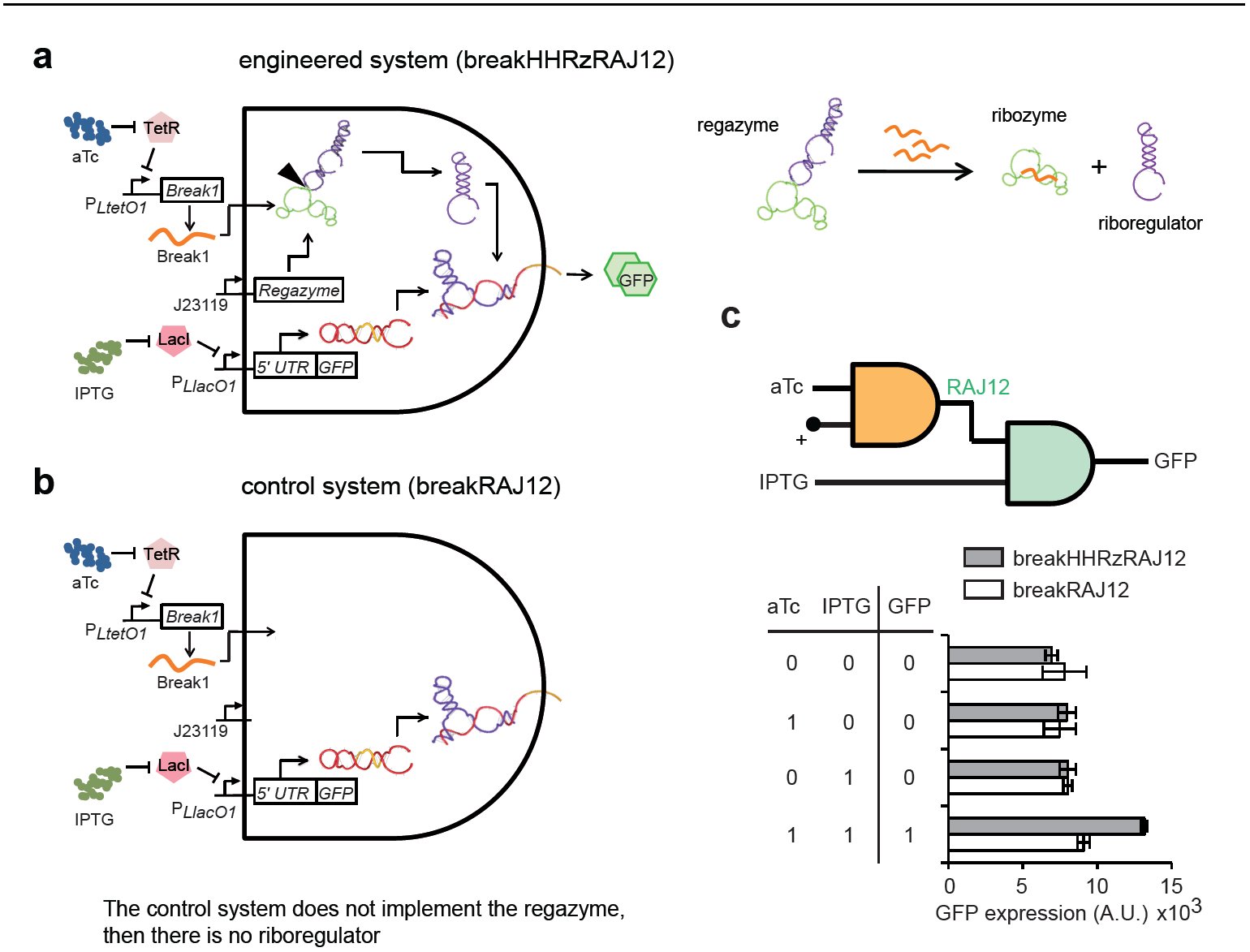
Functional characterization of sRNA-sensing regazyme. (a, b) Schemes of the engineered RNA-based circuit to sense a sRNA and its corresponding control. (c) Digital scheme, associated Truth table, and fluorescence results of regazyme breakHHRzRAJ12 (gray bars), and of its dysfunctional mutant (white bars) for all possible combinations of inducers. Error bars represent standard deviations over replicates.

All together, these results demonstrate the modularity of our designs: *i*) the same sensor module can transduce the signal to different mediators (riboregulators), and *ii*) the same riboregulator (RAJ12 in this case) can be coupled with different sensor modules. We also analyzed experimentally the orthogonality between regazymes. To this end, we constructed new genetic systems based on non-cognate pairs (between riboregulators and 5’ UTRs), in order to test *in vivo* the eventual cross-talk in regulation of gene expression (36). Computational predictions showed no interference between the riboregulators RAJ11, RAJ12 and RR12 (Supplementary Figure 22a), and previous experimental work revealed no apparent activation of sRNA of system RAJ11 on the 5’ UTR of system RAJ12 (18). However, as shown in Supplementary Figure 22b, signaling cross-talk through regazymes can appear (e.g., between RAJ12 and RR12 riboregulatory systems), probably, as a consequence of non-Watson-Crick pairing not covered in the physicochemical model. A further computational design methodology will account for RNA three-dimensional models to better predict RNA-RNA interaction (7) and then engineer RNA circuits with multiple wires. For *N* different sensor modules and *M* orthogonal riboregulators, we could generate, in theory, *NM* regazymes. Importantly, as the output of one regazyme can be the input of another regazyme, we could have at most (*NM*)^*P*^ different implementations of circuits with *P* regazymes, including cascades and feedback loops (Supplementary Figure 23).

### Time-dependent RNA-mediated signal transduction in single cells

To characterize the dynamic response of the designed regazymes at the single cell level, we constructed microfluidics devices according to previous work (36,37). There, single cells were monitored during dozens of cell divisions, using appropriate device geometries to maintain a single layer of cells within the microscope focal plane and a continuous cell growth in exponential phase (Figure 6a, Supplementary Figure 16). Bacterial cells expressing the designed regazymes were loaded into the device, and the composition of the medium was able to incorporate the appropriate time-dependent, small-molecule concentration (25 mM Theo or 100 ng/mL aTc). This way, we can create step functions, pulses or even square waves to force the system. The resulting time series of GFP served to calibrate a mathematical model to further understand the dynamics of the signaling pathway. At this point, we constructed a simplified model, based on first-order kinetics and quasi-steady states assumptions, able to simulate changes in gene expression upon variations in the concentration of the signal molecule (Figure 6b). In essence, the dynamics is modulated by the rates of regazyme cleavage and protein degradation. According to our model, the dynamics can be reduced to an exponential decay with production when the time scale of protein degradation is dominant (Supplementary Figure 21). However, it follows a linear trend in the opposite case. Other parameters such as the cleaved fraction upon signal induction, the effective dissociation constant between the riboregulator and the 5’ UTR, or the gene copy number only affect the stationary level, but not the dynamics. Even though the model can capture small changes in gene expression, the activity could be not discernable in a cellular context when these parameters are suboptimal. Based on our own data, the characteristic times of HHAz (and also HHRz) cleavage and GFP degradation are about 6 and 12 min, respectively (GFP has a degradation tag for riboregulatory devices RAJ12 and RR12). Moreover, the RNA species must also be short lived (about 2 min of half-life) to not exert a significant effect on the dynamics of the systems. In turn, we observed that the response of the system is very fast and without delay, reaching the steady state (criterion of 95%) after 26 min in the case of theoHHAzRAJ12 (Figure 6c). In addition, there is no significant difference in dynamics when inducing with small molecule or sRNA (Figure 6e, see also Supplementary Figures 17, 18). We also analyzed the dynamic response of regazymes theoHHAzRAJ11 (Supplementary Figure 19) and theoHHAzRR12 (Supplementary Figure 20), showing higher dynamic range in these cases and also higher cell-to-cell variability.

**Figure 6.**
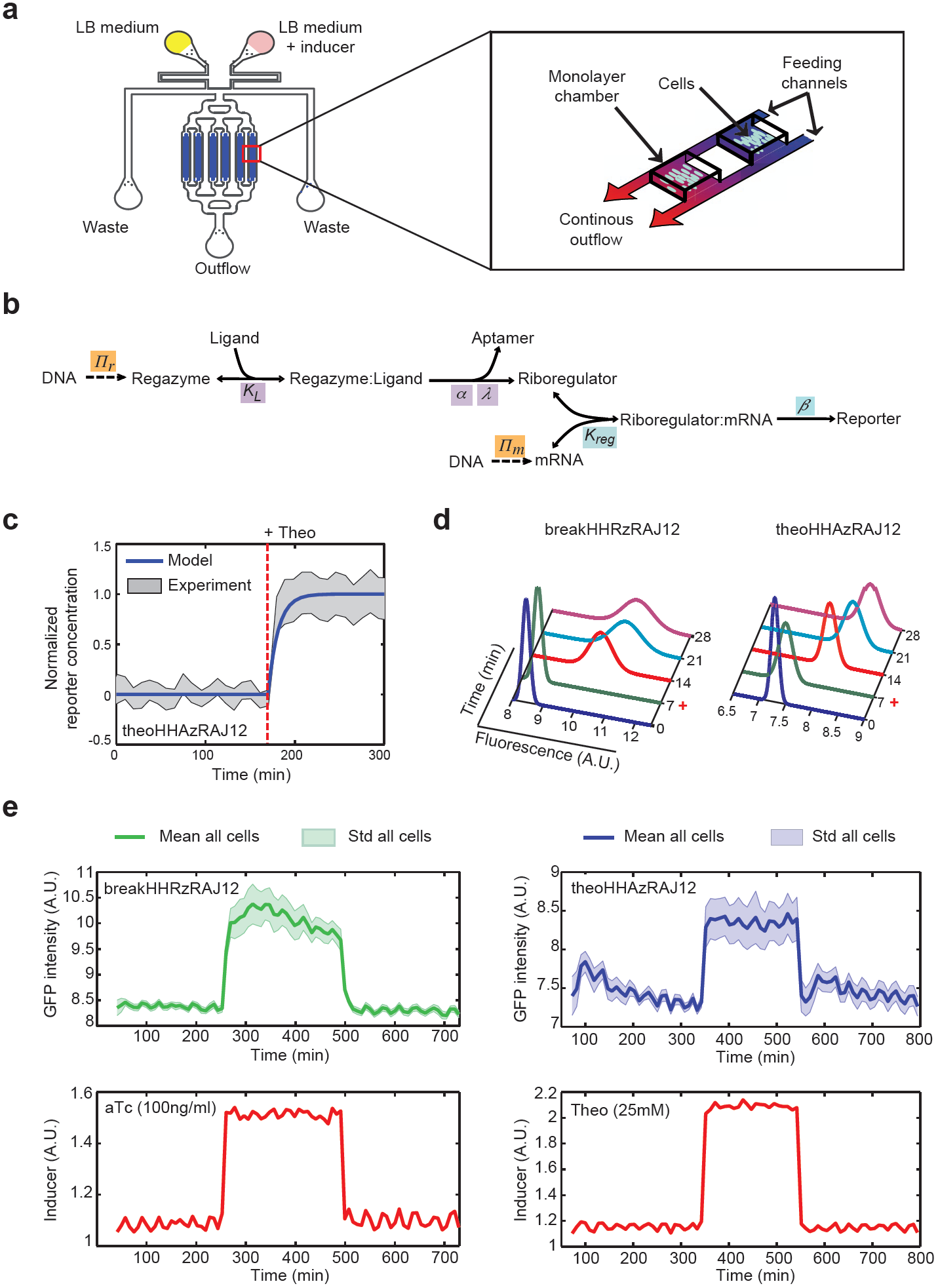
Microfluidics-based single cell analysis of the Theo- and sRNA-sensing regazymes. (a) Scheme of the device used in this work to monitor time-dependent GFP expression in single cells. (b) Scheme of the reactions of the system, together with the relevant parameters, which serve to construct a mathematical model. (c) Prediction of the dynamic response for regazyme theoHHAzRAJ12 (solid line). Shade represents plus/minus one standard deviation from experimental data. (d) Distributions of fluorescence with time across a population of cells and fitted with a Gaussian model. Cells were induced at 7 min. (e) Single cell tracking in one microchamber of fluorescence under a pulse of aTc (100 ng/mL) for breakHHRzRAJ12 or Theo (25 mM) for theoHHAzRAJ12. A constant amount of IPTG (1 mM) was established during the whole experiment. Sulforhodamine B was used to monitor the inducer time-dependent profile.

By collecting all single cell measurements, we can analyze the response of the population and its heterogeneity (quantified as the coefficient of variation –CV– in the state ON). The fluorescence is significantly shifted in all cases after addition of the inducers (Figure 6d). Then, we calculated CV = 0.15 for theoHHAzRAJ12 and CV = 0.17 for breakHHRzRAJ12, revealing a similar heterogeneity in the dynamic response in both cases. However, regazyme theoHHAzRR12 has the wider distribution of the fluorescence (with CV = 0.49), which might be a consequence of a higher dynamic range. Indeed, as the enzymatic degradation machinery limits noise in gene expression (and then cell-to-cell variability) (38), high expression levels can saturate this machinery and then expose the system to noise sources related to growth rate (in the case of theoHHAzRR12, such a saturation is indicated by a slow dynamic response, Supplementary Figure 20). Of importance for further developments, the engineered RNA-mediated signaling pathways are highly responsive to multiple variations in the signal molecule.

## Discussion

We have developed a new kind of synthetic RNA molecules (here termed regazymes) able to transduce signals in a modular way in live cells, introducing the one-to-two-component signal transduction paradigm (Figure 1). The design strategy relies on the hierarchical activation/inactivation of RNA elements with specific function. The design of these cascades of interactions requires, in general, automated design algorithms, which we have implemented thanks to an effective energy model. We have exemplified the strategy by designing several regazymes, engaging different sensor modules responsive to small molecules or sRNAs, and also different mediator modules that work as independent regulatory wires. In turn, each mediator regulates a given reporter gene (here a given 5’ UTR of a common mRNA). The sensor and mediator modules are fused with a transducer module (computationally designed) to form the regazyme molecule. Upon sensing the signal molecule, the aptazyme undergoes a conformational change that activates its self-cleavage activity to release the riboregulator, which is fused in the 3’ end (Figures 2 **and** 3). This riboregulator is then emitted to the cellular context to further activate a downstream *cis*-repressed gene, which may allow the sensing and regulation in different subcellular localizations.

The use of an optimization algorithm, relying on intra- and intermolecular structures and their corresponding free energies as model variables, has allowed us to solve the design problem of the fusion between RNA-based sensors and regulators. However, the computational methods used here to predict free energies and conformational states do not consider three-dimensional contacts neither intermolecular contacts arising in cellular environments (i.e., analysis of the RNA molecules in isolation), which partly limits the predictability of the performance of our designs. By exploring the sequence space, we have identified ideal regions to design functional regazymes without affecting the stability and specificity of the aptamers and riboregulators. The three energetic terms for deriving the objective function have proved effective to capture the functionality of the system (Figure 1). Although we have shown regazymes that sense particular small molecules and sRNAs in live cells, there are more possibilities to widen the scope of their use with the development of new aptazymes. For example, different RNA aptamers sensing chemicals like arginine, nucleotides like flavin, or proteins like HIV-1 Rev (39) encourage the design of the corresponding regazymes by combining computational and experimental screening techniques. This would allow the creation of systems to diagnose the physiological state of a cell, e.g., if it has a new metabolic route in action, if it grows faster than normal, or if it is infected by a virus. In addition, we have demonstrated the versatility of our regazymes by drawing on different riboregulators, with the toehold region in the 5’ end, as mediators. This modularity has the potential to increase the fan-in or fan-out of the system (i.e., the number of inputs and outputs), by disposing in battery regazymes with various sensor domains (signal integration) or with various mediator domains (signal spread). This will result in RNA-mediated signaling hubs. The decoupling of the activities of the aptazyme and riboregulator, besides allowing the straightforward creation of novel systems in a combinatorial way, can enhance the predictability of the dynamic response and the quality of the transmitted information. Similarly, a cleavage-based decoupling of the 5’ UTR sequence from upstream regulatory elements (mainly a promoter) was recently proposed for a protein expression context (40).

Nucleotide-level energy models, such as the one presented in this work, can provide the underlying explanations for molecular interactions, which can then be used in the construction of higher-level biological systems (7). Our study has shown the capability of RNA to mediate in signaling pathways in a cellular context similar to proteins. However, engineering signal transduction with RNA bypasses the challenges related to membrane localization, complex assembly, competition, and limitation of interaction modules regular to protein-based two-component systems (2). Then, the use of regazymes, as pre-engineered modules, may allow devising more complex RNA circuits, at the same time they illustrate new ways to engineer arbitrary complexity. We have shown that a regazyme can be used to engineer multi-input AND logic gates, through the combination of transcriptional and post-transcriptional regulations (**Figures 4 and 5**). Yet, nothing prevents, in theory, using complementary riboregulatory elements such as RNA-binding/processing proteins (41) or translation-transcription control adaptors (42) to generate feedback or feedforward loops. Therefore, a regazyme could serve as an information transmission platform to control over network connections in live cells, which promises future synthetic biology developments, such as metabolic control and rewiring (9).

In addition, we have measured the dynamic response of the catalytic activity of our designed regazymes in cellular RNA extracts (Figures 2 **and** 3), obtaining comparable values than those previously measured *in vitro* with the same ribozyme core (11,32). These results have revealed a fast cleavage rate, which is within the same time-scale as protein modifications in synthetic systems (5). Moreover, we have observed higher heterogeneity of the dynamic response (from cell to cell) for the sRNA-sensing regazyme, pointing out an effect of heterogeneous gene expression or even RNA-RNA interaction across cells. Using a mathematical model, we have also analyzed the impact on the dynamics of key parameters governing this RNA-mediated signaling pathway. We have further characterized in detail the dynamic response of the riboregulated gene by monitoring a single cell expressing a regazyme using a microfluidics platform (Figure 6). Systems with higher dynamic range also exhibit higher heterogeneity in their dynamic response (from cell to cell). Our results have shown, in agreement with the catalytic assays, a fast activation of protein expression.

The modularity of our approach, where the sensing and mediator modules can be designed independently through computational design of a proper interface, would allow us to replace the mediator module to control eukaryotic gene expression. Perhaps the best strategy would consist in expressing the regazyme only in the nucleus, using a suitable promoter (e.g., a promoter transcribed by RNA polymerase III). One example would consist in exploiting as mediator module a sRNA able to guide the CRISPR-associated catalytically inactive dCas9 protein to block transcription initiation or elongation (recently proposed as CRISPR interference in both prokaryotes and eukaryotes (43,44)). This RNA-guided DNA targeting would allow the addition of a transcription regulation layer into the RNA-mediated signal transduction. Another example of application in eukaryotic cells would consist in fusing micro RNAs (miRNAs) rather than bacterial riboregulators with sensor domains. There, an aptazyme engineered in the 3’ UTR of the primary miRNA controlling the poly(A) tail could serve to link the mediator to the signal molecule (12). In conclusion, we envision our computationally designed regazymes as a start point for engineering more sophisticated RNA-mediated signaling with particular applications. This would allow the synthetic biology community to use RNA devices to incorporate new signaling functions into cells or rewire natural protein signaling pathways (45).

## Acknowledgements

We thank Mr. W. Rostain for proofreading the manuscript.

## Supplementary Information

### Supplementary Materials and Methods

#### Plasmids construction

To express and characterize the RNA-based signal transduction, we have generated a modified plasmid vector, pSTC2 containing a pSC101m origin of replication (a mutated pSC101 ori giving a high copy number) and a kanamycin resistance selection marker (Supplementary Figure 1). The pSTC2 vector is based on our previously reported vector pSTC1^1^ by removing the mRFP coding sequence and tagging the carboxyl terminus of the superfolder GFP (sfGFP)^2^ with the *ssr*A degradation tag (<u>AS</u>AANDENYALAA)^3^ by polymerase chain reaction (PCR). The underlined AS dipeptide coding sequence corresponds to an NheI restriction site and acts as a linker, while the *ssr*A tag targets proteins to the ClpXP degradation pathway, significantly increasing their degradation rates and therefore dynamic behaviors^3^,^4^.

For engineering our gene cassettes, the pSTC2 vector was made so that independent promoters could drive the expression of the regazyme and the corresponding mRNA. We used the inducible promoters P_LlacO1_ (regulated by LacI and modulated externally by the chemical inhibitor isopropyl-β-D-thiogalactopyranoside, IPTG) and P_LtetO1_ (regulated by TetR and modulated externally by the chemical inhibitor anhydrotetracycline, aTc)^5^. Note that both promoters were placed in opposite directions to avoid transcriptional interference^1^. The regazymes were under the control of the promoter P_LtetO1_, and the reporter mRNAs (with the *cis*-regulating elements) were under the control of the promoter P_LlacO1_. For the sRNA-sensing regazyme (breakHHRzRAJ12), the signal sRNA (break1) was placed under control of the promoter P_LtetO1_ and the regazyme was under the control of the constitutive promoter J23119. The sequences of the regazymes and the corresponding 5’ UTRs of the targeted mRNAs are presented in the Supplementary Tables 1-4. The list of plasmids used in this study is given in the Supplementary Table 6. All plasmid manipulations were performed using standard molecular biology techniques^6^. All enzymes used for plasmid digestions were from Thermo Scientific, USA. All oligonucleotides were synthesized from Integrated DNA Technologies, USA. The different RNA devices (from the terminator of the regazyme to the 5’ UTR of the mRNA, see Supplementary Figure 2) were chemically synthesized and cloned in plasmid pIDTSMART (pUC replication origin, ampicillin resistance marker) and then subcloned into pSTC1 or pSTC2.

#### PCR-based mutagenesis

Dysfunctional regazymes (both core catalytic mutations and theophylline binding activity mutations in the aptamers, see Supplementary Figure 7) were constructed using PCR-based site-directed mutagenesis with Phusion high fidelity DNA polymerase (Thermo Scientific, USA), followed by template digestion with DpnI (Thermo Scientific, USA) for 1 h at 37 μC. The final products were transformed into chemical competent *E. coli* cells. Mutations were further confirmed by plasmid sequencing (GATC ®, Germany). The primers for PCR-based mutagenesis in the theophylline aptamer were 5’-aatccaggacacccgcccagggcctttcggc-3’ and 5’-gccgaaaggccctgggcgggtgtcctggatt-3’ for theoHHAzRAJ12, 5’-aatccaggacacccgcccagggcctttcggc-3’ and 5’-gccgaaaggccctgggcgggtgtcctggatt-3’ for theoHHAzRAJ11, and finally 5’-gccgaaaggccctgggcgggtgtcctggatt-3’ and 5’-aatccaggacacccgcccagggcctttcggc-3’ for theoHHAzRR12. In addition, for breakHHRzRAJ12 we used the following primers 5’-ctcgtcgatccctccctatcagtgatagagattg-3’ and 5’-caatctctatcactgatagggagggatcgacgag-3’ to excise the regazyme sequence in order to have a dysfunctional system (see Supplementary Figure 13c).

#### Strains, reagents and cell cultures

Strains used in this study are listed in the Supplementary Table 6. *E. coli* strain DH5α (Invitrogen, USA) was used for plasmid construction purposes as described in the protocol^6^. Characterization experiments were performed in *E. coli* K-12 JS006 cells (MG1655 Δ *araC* Δ*lacI*)^7^, and/or in *E. coli* K-12 MG1655Z1 (or simply MGZ1) cells (MG1655 *lacI*^+^ *tetR*^+^ *araC*^+^ Sp^R^)^8^ for control over the promoters P_LlacO1_ and P_LtetO1_. Cells were grown aerobically in Luria-Bertani (LB) broth or in a modified M9 minimum medium, prepared with M9 salts (Sigma, Germany), glycerol (0.8%, vol/vol) as only carbon source, CaCl_2_ (100 μM), MgSO_4_ (2 mM), and FeSO_4_ (100 μM). Cultures were grown overnight at 37 μC and at 225 rpm from single-colony isolates before being diluted for *in vivo* characterization. When appropriate, kanamycin concentration was 50 μg/mL and ampicillin concentration was 100 μg/mL. In the case of MG1655Z1 cells, 1 mM IPTG (Thermo Scientific, USA) was used for full activation of promoter P_LlacO1_ when needed, and 100 ng/mL aTc (Sigma, Germany) was used for full activation of promoter P_LtetO1_. 4 mM theophylline (Sigma, Germany) and 0.5 mM thiamine hydrochloride (Sigma, Germany) were used for general characterization of the small molecule sensing regazymes in the fluorometer. We also used gradients of aTc, theophylline, and thiamine to perform dose-dependent assays. For microfluidic cell cultures, cells were grown aerobically in fresh LB broth or in LB supplemented with 0.05% sulforhodamine B (Sigma, Germany) and (i) 25 mM theophylline or (ii) 100 ng/mL aTc.

#### Quantification of *in vivo* fluorescent protein synthesis

Cells were grown overnight in 5 mL of LB medium, and were refreshed in culture tubes with LB medium in order to reach stationary phase. Cells were then diluted 1:200 in 200 μL of M9 minimal medium in each well of the plate (Custom Corning Costar 96 well microplate, black transparent bottom with lid). The plate was incubated in an Infinite F500 multi-well fluorometer (TECAN, Switzerland) at 37 °C with shaking (orbital mode, frequency of 33 rpm, 2 mm of amplitude). It was assayed with an automatic repeating protocol of absorbance measurements (600 nm absorbance filter) and fluorescence measurements (480/20 nm excitation filter - 530/25 nm emission filter for sfGFP) every 15 min. All samples were present in triplicate on the plate. Each measurement was repeated on independent days to verify reproducibility. All data analyses were done using values harvested when cells were in exponential growth phase (OD_600_ between 0.1 and 0.6). Growth rates were calculated as the slope of a linear regression between the values of ln(OD_600_) and time. The normalized fluorescence was obtained by two methods: (*i*) as the slope of the linear regression between the values of absolute fluorescence (*F*) and OD_600_, and (*ii*) as the ratio between absolute fluorescence and OD_600_ (i.e., 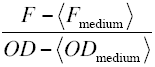, where *brakets* denote average per samples). The normalized fluorescence of plain cells (transformed with a plasmid without GFP) was also considered for background subtraction when appropriate, and then obtaining the stationary protein expression value (magnitude per cell).

#### Quantification of *in vivo* catalytic activity

To *in vivo* quantify the catalytic activity of the two versions of the RAJ12-based regazyme that sense theophylline (theoHHAzRAJ12) and an effector RNA (breakHHRzRAJ12), we transformed *E. coli* (strain T7 Express in the case of theoHHAzRAJ12 and strain MG1655Z1 in the case of breakHHRzRAJ12) with the appropriate plasmids (pSCKtheoRAJ12 for theophylline-induced cleavage, and pUAbreak12 and pSCKbreak12 for sRNA-induced cleavage). Overnight cultures were grown at 28 °C by inoculating LB liquid media containing 50 μg/mL kanamycin (theoHHAzRAJ12) or 50 μg/mL kanamycin and 50 μg/mL spectinomycin (breakHHRzRAJ12) with three different colonies in each case. These overnight cultures were used to inoculate new liquid cultures with 50 mL of the corresponding media at 0.1 OD_600_ and grown at 37 °C to reach 0.6 OD_600_. At this point each culture was split into three aliquots of 15 mL, and theophylline or aTc was added when appropriate to obtain cultures with 0, 0.4 and 4 mM theophylline or 0, 10 and 100 ng/mL aTc. Incubation was continued at 37 °C with shaking and 2 mL aliquots were taken at 0, 2, 4, 8, 16 and 32 min time points. Bacteria in the 2-mL aliquots were quickly pelleted by centrifuging for 2 min at 13,000 rpm and re-suspended in 50 μL of TE (10 mM Tris-HCl, pH 8.0, 1 mM EDTA). Bacteria were broken by adding 50 μL of a 1:1 phenol:chloroform (pH 8.0) mix and vortexing thoroughly. Bacterial RNA from each sample was recovered in the aqueous phase by centrifuging for 5 min at 13,000 rpm, re-extracted with 50 μL chloroform.

Processing extent of regazyme in each aliquot was analyzed by northern blot hybridization using a complementary [^32^P]-labelled RNA probe after separating the different RNA samples by denaturing polyacrylamide gel electrophoresis (PAGE). 20 μL of the RNA preparations were mixed with one volume of formamide loading buffer (98% formamide, 10 mM Tris-HCl, pH 8.0, 1 mM EDTA, 0.0025% bromophenol blue, and 0.0025% xylene cyanol), denatured for 1.5 min at 95 °C and snap cooled on ice. After this treatment, samples were separated by PAGE in 5% polyacrylamide (37.5:1 acrylamide:N,N’- methylenebisacrylamide) gels of 140 × 130 × 2 mm including 8 M urea and TBE buffer (89 mM Tris, 89 mM boric acid, 2 mM EDTA) for 1.5 h at 200 V. Gels were stained with ethidium bromide, photographed under UV light and the RNAs were electroblotted to positively charged nylon membranes (Nytran SPC; Whatman, USA) and cross-linked by irradiation with 1.2 J/cm^2^ UV light (Vilber Lourmat). Membranes were hybridized overnight at 70 °C in 50% formamide, 0.1% Ficoll, 0.1% polyvinylpyrrolidone, 100 ng/mL salmon sperm DNA, 1% SDS, 0.75 M NaCl, 75 mM sodium citrate, pH 7.0, and 10^5^ cpm/mL of the [^32^P]-labelled complementary RNA probe. Membranes were washed three times for 10 min with 2 × SSC (SSC is 150 mM NaCl, 15 mM sodium citrate, pH 7.0), 0.1% SDS at room temperature and once for 15 min at 55 °C with 0.1 × SSC, 0.1% SDS. Membranes were imaged by autoradiography and the hybridization signals quantified by phosphorimetry (Fujifilm FLA-5100, Japan).

To produce the [^32^P]-labelled RNA probe in order to quantify regazyme processing, we firstly amplified by PCR a fragment of the regazyme cDNA using primers 5’- TGGCGCTGCCTTCGTACATCC-3’ and 5’-ACAGAAAAGCCCGCCTTTCGA-3’. The probe corresponds to the reverse complement of regazyme theoHHAzRAJ12 and was used to detect both the regazymes and cleaved products of systems theoHHAzRAJ12 and breakHHRzRAJ12. This cDNA was cloned in the appropriate orientation into a pUC18 (L08752.1)-derived plasmid flanked by a bacteriophage T3 RNA polymerase promoter and an XbaI restriction site. The *in vitro* transcription reaction consisted of 1 μg of the XbaI-linearized plasmid, 2 mM each ATP, CTP and GTP, 70 μCi of [α-^32^P] UTP (800 Ci/mmol), 40 mM Tris-HCl (pH 8.0), 6 mM MgCl_2_, 20 mM DTT, 2 mM spermidine, 20 U RNase inhibitor (Ribolock, Thermo Scientific, USA), 0.1 U yeast inorganic pyrophosphatase (Thermo Scientific, USA) and 50 U of T3 RNA polymerase (Epicentre, USA) in a final volume of 20 μ L. The reaction was incubated for 2 h at 37 °C. Then, 10 U of DNase I (Thermo Scientific, USA) were added and incubated at 37 °C continued for 10 min. The probe was finally purified by chromatography using a Sephadex G-50 spin column (Mini Quick Spin Column, Roche Applied Science) and quantified by Cerenkov.

#### Quantification of *in vitro* self-cleavage activity

We cloned the regazymes theoHHAzRAJ12 and breakHHRzRAJ12 (without transcription terminators) into new plasmids (pUC-like) under the control of a T3 promoter and by adding the sequences GGGAT in the 5’ end and ATCTCTAG (this is XbaI restriction site) in the 3’ end. Linearization of the plasmids was done with XbaI, followed by purification with silica-based columns (ZYMO). Reactions of *in vitro* transcription were carried out without and with (4 mM) theophylline in case of theoHHAzRAJ12, and without and with (3 μM) the DNA oligo Break1 in case of breakHHRzRAJ12. 100 ng and 70 ng of plamids having regazymes theoHHAzRAJ12 and breakHHRzRAJ12 were used for the reactions, respectively. Reaction in a final volume of 20 μL was done with 2 μL transcriptase buffer 10x, 0.4 μL 0.5 M DTT, 1 μL 10 mM NTPs, 0.5 μL RNase inhibitor Ribolock (Thermo Scientific, 40 U/μL), 1 μL inorganic pyrophosphatase (Thermo Scientific, 0.1 U/μL), 1 μL T3 RNA polymerase (Roche, 20 U/μL). We incubated for 30 min at 37 μC. Then we added 20 μL (1 vol.) formamide buffer to stop the reaction. Samples were then heated 1.5 min at 95 °C, followed by storage on ice. To load the gel (5% PAGE, 8 M urea, TBE 1x), we mixed half of the resulting sample (10 μL) with the appropriate buffer (10 μL). The conditions were 200 V and 1.5 h. We used the Thermo Scientific RiboRuler Low Range RNA Ladder.

For the *in vitro* time-course of self-cleavage assay, plasmid containing regazyme theoHHAzRAJ12 was linearized with XbaI and transcribed with T3 RNA polymerase in the presence of high NTPs concentration (2 mM each) to sequester free Mg^2+^ in the reaction and inhibit regazyme self-cleavage. Transcription products were separated with denaturing PAGE, and the uncleaved regazyme eluted from the gel by diffusion in the presence of 10 mM EDTA. Regazyme was precipitated with ethanol and finally resuspended in 50 mM Tris-HCl, pH 8.0, 1 mM EDTA. A time-course self-cleavage experiment in the presence or not of 4 mM theophylline was performed. Both reactions were started by adding MgCl_2_ to 5 mM and aliquots taken at 0, 1, 2, 4, 8 and 16 min. Reactions were stopped using a buffer including formamide and EDTA. Finally, reaction products in the different aliquots were separated with denaturing PAGE in the presence of 8 M urea. The gel was stained with ethidium bromide and the bands corresponding to the uncleaved regazyme and the cleavage products quantified through the fluorescent emission under ultraviolet irradiation. A similar procedure was carried out to analyze the self-cleavage activity of regazyme breakHHRzRAJ12.

#### Microfluidics device construction

To understand the dynamic regulation of our regazyme devices, we have examined *in vivo* gene expression with single-cell, time-lapse fluorescence microscopy using a microfluidics device (Supplementary Figure 16a). This was designed to support monolayer growth of *E.coli* cells under constant nutrient flow. By coupling with cell tracking and fluorescence measurement, our microfluidics device allows us to generate fluorescence trajectories for single cells. The design of the microfluidics device was adapted from the previous one reported by Hasty and coworkers^7^,^9^. In brief, the *E. coli* cells were loaded from the cell inlet while keeping the media inlet at sufficiently high pressure to avoid contamination. Cells were loaded into the microchambers by manually applying pressure pulses to the syringe lines to induce a momentary flow change. After cell loading, the flow was then reversed to allow cells receiving fresh media with 0.075% Tween20 which prevented cells from adhering to the main channels and waste outlets. The microfluidics device contained three parallel channels with 10 μm height. Each channels contained two subchannels with microchambers (1 μm height) in the middle of them, which allowed the out growing cells been washed away by the flow. The width of the parallel chamber was limited to 30 μm to avoid the risk of the PDMS structural collapse of the chamber ceilings. For optimal *E. coli* growth, the microfluidics chip temperature was typically maintained at 37 μC by external tempcontrol system (PECON, Germany). For on-chip induction experiments, we used two media inlets (for two different input media, with and without chemical inducer), which directed the flow to the cells from one to another by means of pressure changes. For induction flow (medium with inducer), the media input speed was kept at 500 μL/h, and for relaxation flow (medium without inducer) was kept at 10 μL/h.

The design of the microfluidics device was performed in AUTOCAD software (AUTODESK, USA), and the printed wafer was fabricated by Veeco Instrument GmbH (Veeco, Germany). Replica molds were created *in house* from master molds by mixing polydimethylsiloxane (PDMS)/Sylgard 184 (Dow Corning, USA) in a 10:1 ratio of elastomer base vs. curing agent. The molds were degassed by briefly centrifuge at 4000 rpm for 5 min, followed by degassing in a vacuum desiccator at −1 a.t.m. for 30 min, and curing in place over the master at 80 μC for 2 h. After removal of the PDMS monolith, chips were sectioned, bored at the fluidic ports in a clean culture hood, and then bonded to clean coverslips (Corning, USA) by exposure to O_2_ plasma at 30 W for 30 s in a plasma asher (ACE-1, GaLa Instrumente, Germany). The bonded chips were cured in 80 μC incubator for at least 2 days before experiments.

#### Time-lapse microscopy and image analysis

All images were acquired using Zeiss Axio Observer Z1 microscopy (Zeiss, Germany), comprising a Pln Apo 100X/1.4 Oil Ph3 DICIII objective outfitted with fluorescence excitation and emission filter wheels (including 38HE GFP RL, 46HE YFP RL, 47HE CFP RL, and 64HE RFP RL). The microscope resolution was 0.24 μm with Optovariation 1.6X, resulting total magnification 1600X for both bright field and fluorescent images. The microscopy equipped with an X-Cite Series 120 fluorescent lamp (EXFO, illumination) and a HAMAMATSU EM-CCD C9100 digital camera (Hamamatsu Photonics, Japan) automated by a commercial application software (AxioVs 40 V4.8.1.0, Zeiss, Germany).

In each experiment, the microfluidics device mentioned above was mounted to the stage and loaded with *E. coli* cells. Trapped cells were allowed to grow overnight with normal LB medium flow at 500 μL/h. During exponential growth of the monolayer colony, images were collected at 100X magnification in the phase contrast every 30 s and GFP or RFP fluorescence channels every 7 min (GFP channel for monitoring the gene expression, RFP channel for monitoring the inducer diffusion) over a period of around 24 h (Supplementary Figures 16b and 16c). In each experimental run, we have chosen four different cell microchambers to follow the gene expression dynamics. Focus was predefined by focus setting with contrast-based autofocus algorithms. Images were extracted and movies were generated by *in house* processing based MATLAB (MathWorks, USA) for both phase contrast and fluorescent frames respectively. Briefly, inverted phase contrast movies were used for threshold settings, followed by generating the binary threshold phase-contrast movies. To compensate the binary movies for minor movements of different frames due to the microscopy, large objects were found in order to track the stability. After this correction, the resulting binary image was processed with morphological distance transformation and watershed segmentation. Cells were tracked by defining a cell-to-cell distance matrix and the cell lineages were reconstructed. Finally the fluorescence level of each cell in each fluorescence frame was extracted.

#### Optimization algorithm

We developed an optimization algorithm to design regazymes provided the sequences of a given aptazyme and a riboregulator. On the one hand, the aptazyme responds to its ligand to cleave the RNA sequence at a given point^10^. On the other hand, the riboregulator is able to activate protein expression by inducing a conformational change in the 5’ UTR of the mRNA. The sequence of a regazyme is composed of prefix and suffix sequences flanking the aptazyme followed by the riboregulator (Supplementary Figure 6a). These sequences constitute part of the transducer module. The aptazyme and riboregulator sequences are kept fixed. The only premise for the design is that the riboregulator needs to have the seed region in its 5’ tail. Hence, the prefix and suffix are designed to get the seed paired within the structure of the full regazyme (state OFF), and unpaired (and then exposed to the solvent for RNA-RNA interaction) within the resulting structure after cleavage induced by the ligand (state ON). These sequences are totally variable in nucleotide composition and length. Starting from random sequences for the prefix and suffix, the algorithm implements a heuristic optimization based on Monte Carlo Simulated Annealing (Supplementary Figure 6b)^10^. At each step, random mutations consisting in replacements, additions, or deletions are applied to evolve the sequence, and then selected with an objective function (Δ*G*_score_). This way, we constructed the minimization problem

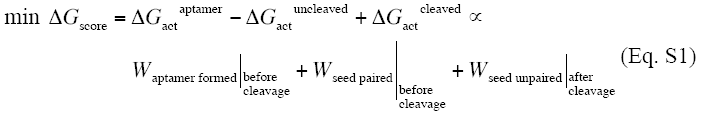

where *W* is the thermodynamic work required to get the aptamer formed to sense the ligand or get the seed unpaired or paired, and it is assumed to be proportional to the activation free energy (Δ*G*_act_). We balanced equally these states. Denoting by Λ and Γ_0_ the structures of the aptamer and seed within the regazyme before cleavage, and by Γ the structure of the seed after cleavage, we can calculate these works by

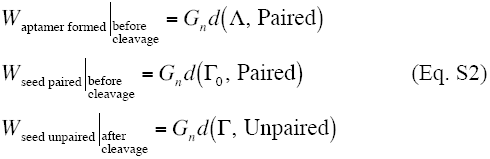

where *G*_*n*_ is the average free energy contribution per nucleotide (here we consider *G*_*n*_ = 1.28 Kcal/mol), and *d* is the Hamming distance between two secondary structures.

Typically, the convergence of the algorithm is fast, obtaining sequences with **Δ***G*_score_ = 0 in some seconds or minutes. We used the Vienna RNA package with default parameters^11^. Our systems are based on conformational changes, but algorithms for multi-state RNA design are scarce. Our approach tackles this problem, allowing sequence and structure specifications, and exploiting RNA folding algorithms such as Vienna RNA. In this work, we just focused on the computational design of the transducer module, but nothing would prevent a full design of the molecule, including the riboregulator. A strategy of nesting design processes (one for the riboregulator and another for the regazyme) enhances the convergence of the corresponding algorithms, as well as makes the designs more modular. To this end, our approach has the advantage of leaving unconstrained the sequence length. However, our approach has the limitation of just using 2D structure to model RNA conformational change and catalysis. Certainly, this type of mechanisms could involve pseudoknot interactions and even non-canonical base pairs, for which 3D models could better capture the interaction and processing features. Nevertheless, although the ribozyme has tertiary contacts, the exposition or blockage of the seed region in our design is governed by secondary structure. In addition, our model does not take into account kinetic binding effects, which might have an impact on the designs.

#### Mathematical model

The full set of biochemical reactions (constants in brackets) of the system theoHHAzRAJ12 (small molecule sensing) is

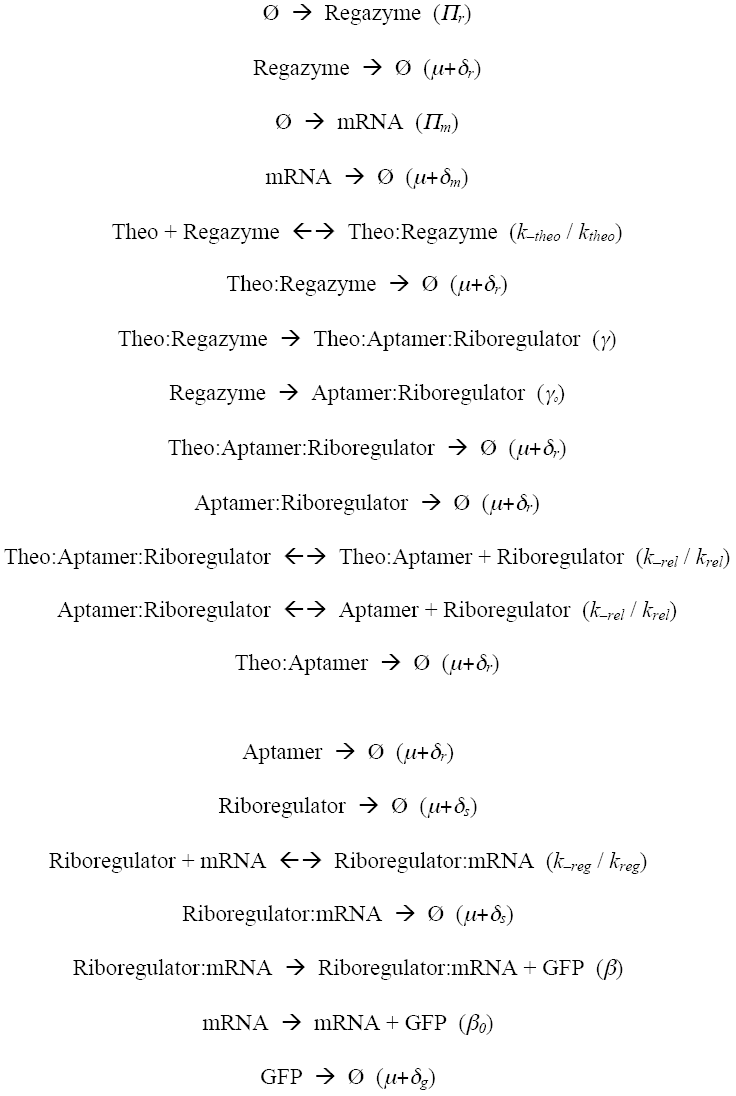

Clearly, this model can be easily rewritten in case of sRNA sensing. Then, to quantitatively model the protein synthesis in the cells, we could construct a system of differential equations based on those reactions. However, due to the lack of reliable values for many of the parameters, we decided to take a quasi-steady state approach.

We first assumed that the global and intracellular concentration of external inducers is the same, and that they bind very fast (relative to other time scales in the system) to their target molecules (i.e., IPTG to LacI, aTc to TetR, and theophylline –Theo– to RNA aptamer). Therefore, the total concentrations of RNAs are taken in quasi-steady state, given by

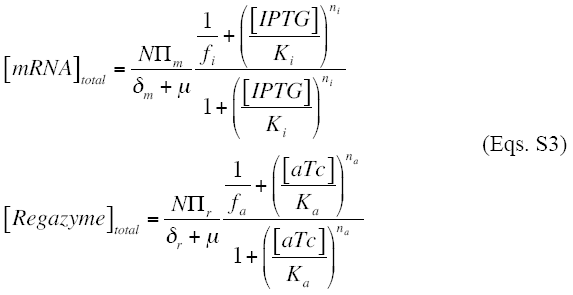

where IPTG and aTc can vary with time. *N* is the plasmid copy number. *Π*_x_ (x = *m* or *r*) is the maximal transcription rate of the transcript (mRNA or regazyme), and *δ*_x_ (x = *m* or *r*) the corresponding degradation rate (mRNA or regazyme). The growth rate of the cells is *μ*. In addition, *K*_x_ (x = *i* or *a*) is the effective regulatory constant of the inducer (IPTG or aTc) that inhibits the repressor action (of LacI or TetR), and *n*_x_ (x = *i* or *a*) the effective Hill coefficient (for IPTG or aTc).

Furthermore, because the concentration of theophylline is much higher than the one of regazyme, we can write

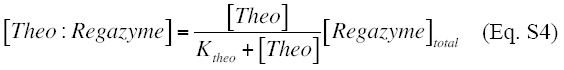

where *K*_*theo*_ is the effective dissociation constant *in vivo* between the aptazyme and theophylline (i.e., *K*_*theo*_ = *k*_–*theo*_ / *k*_*theo*_). The concentration of free regazyme is then [*Regazyme*] = [*Regazyme*]_*total*_ – [*Theo:Regazyme*].

To simplify the model of the aptazyme cleavage, we introduce the following term

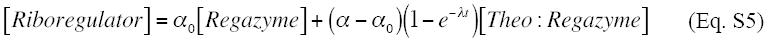

where *α*_0_ is the fraction of cleavage in absence of any ligand and *α* the maximal fraction in presence of theophylline. Moreover, *λ* is rate at which the catalytic reaction takes place (it is related to *γ*). Previous to the addition of theophylline, we can write [*Regazyme*] = [*Regazyme*]_*total*_. After the addition of theophylline in high amount, we can consider [*Theo:Regazyme*] = [*Regazyme*]_*total*_.

Once the riboregulator is released, it can interact with its mRNA target. We assume this reaction is faster than that of cleavage and does not introduce any delay. Hence, the concentration of the resulting complex is

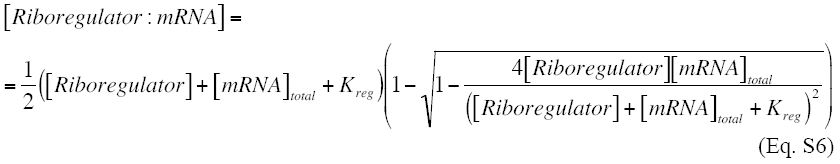

where *K*_*reg*_ is the effective dissociation constant *in vivo* between the riboregulator and mRNA (i.e., *K*_*reg*_ = *k*_–*reg*_ / *k*_*reg*_). The concentration of free mRNA is then [*mRNA*] = [*mRNA*]_*total*_ – [*Riboregulator:mRNA*]. Therefore, the synthesis of GFP is governed by

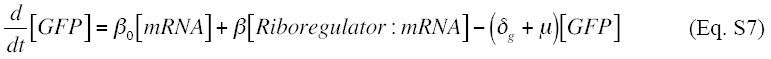

where *β*_0_ is the translation rate in absence of riboregulator and *β* the rate in presence of it. Finally, *δ*_*g*_ is the first-order protein degradation rate. [*GFP*]_0_ = 0 could be taken as initial condition *in vitro*, but because the system is expressed in a cellular context we should take [*GFP*]_0_ = *β*[*Riboregulator*: *mRNA*] /(*δ*_*g*_ + *μ*) calculated with [*Riboregulator*] = *α*_0_ *M*.

#### Mathematical analysis of the dynamic response

To analyze the dynamic response of the regazyme system, we assume that the only inducer with time dependence is theophylline and that the total concentrations of mRNA and regazyme are in steady state. Thus, from Eqs. (S3) and considering *δ*_*m*_ = *δ*_*r*_, we have

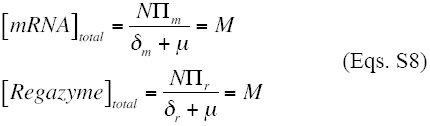

where *M* ≈ 3000 nM for a particular set of parameter values (Supplementary Table 5). We also assume that [*Theo*] ≫ *K*_*theo*_ to write Eq. (S5) as

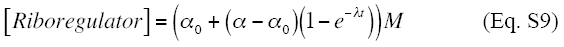

In case of producing directly a riboregulator from promoter P_LtetO1_, we would have [*Riboregulator*] = *M*. In addition, because the *cis*-repression is very efficient^1^ and GFP has a degradation tag, we can take *β*_0_ ≪ *β* and *δ*_*g*_ >> *μ*. Finally, we can write the time dependence of GFP as

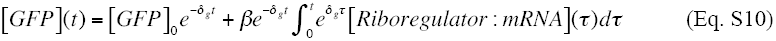

to be integrated numerically in combination with Eqs (S6) and (S9).

## Supplementary Figure Legends

**Supplementary Figure 1:**
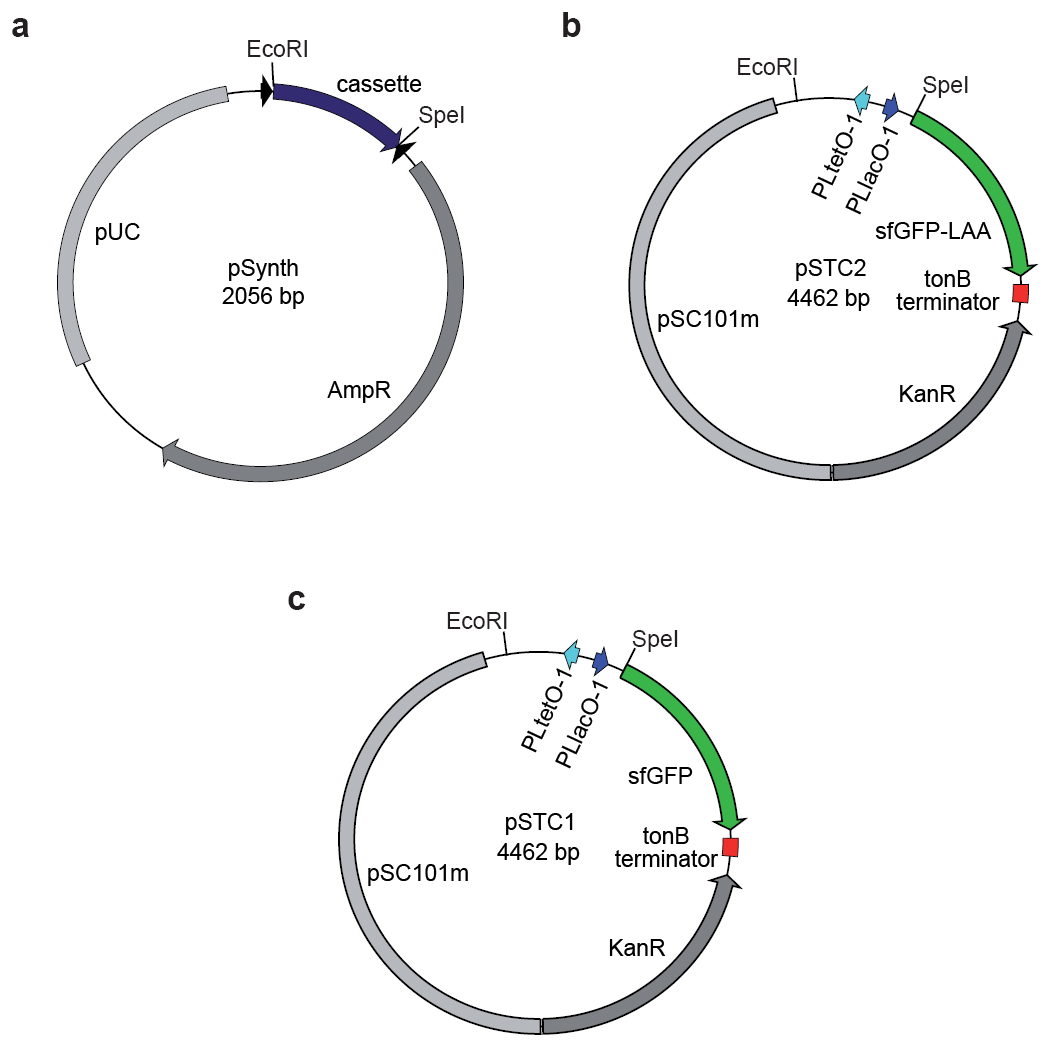
Maps of the plasmid vectors used in this work for expressing the designed regazymes. (**a**) Plasmid vector map for gene synthesis of the regazymes used in this study. This vector contains a high copy number pUC replication origin and an ampicillin resistance marker. (**b**) pSTC2 vector map. (**c**) pSTC1 vector map. Gene cassettes from pSynth can be cloned into these vectors by using restriction digestion with EcoRI and SpeI. Except otherwise indicated, regazymes were under the control of the inducible promoter P_LtetO1_, and the reporter mRNAs (with the *cis*-regulating elements) were under the control of the inducible promoter P_LlacO1_. Vectors pSTC1 and pSTC2 contain a high copy number pSC101m replication origin (mutated version of pSC101) and a kanamycin resistance marker. Maps were constructed with the free software Savvy (http://bioinformatics.org/savvy/).

**Supplementary Figure 2:**
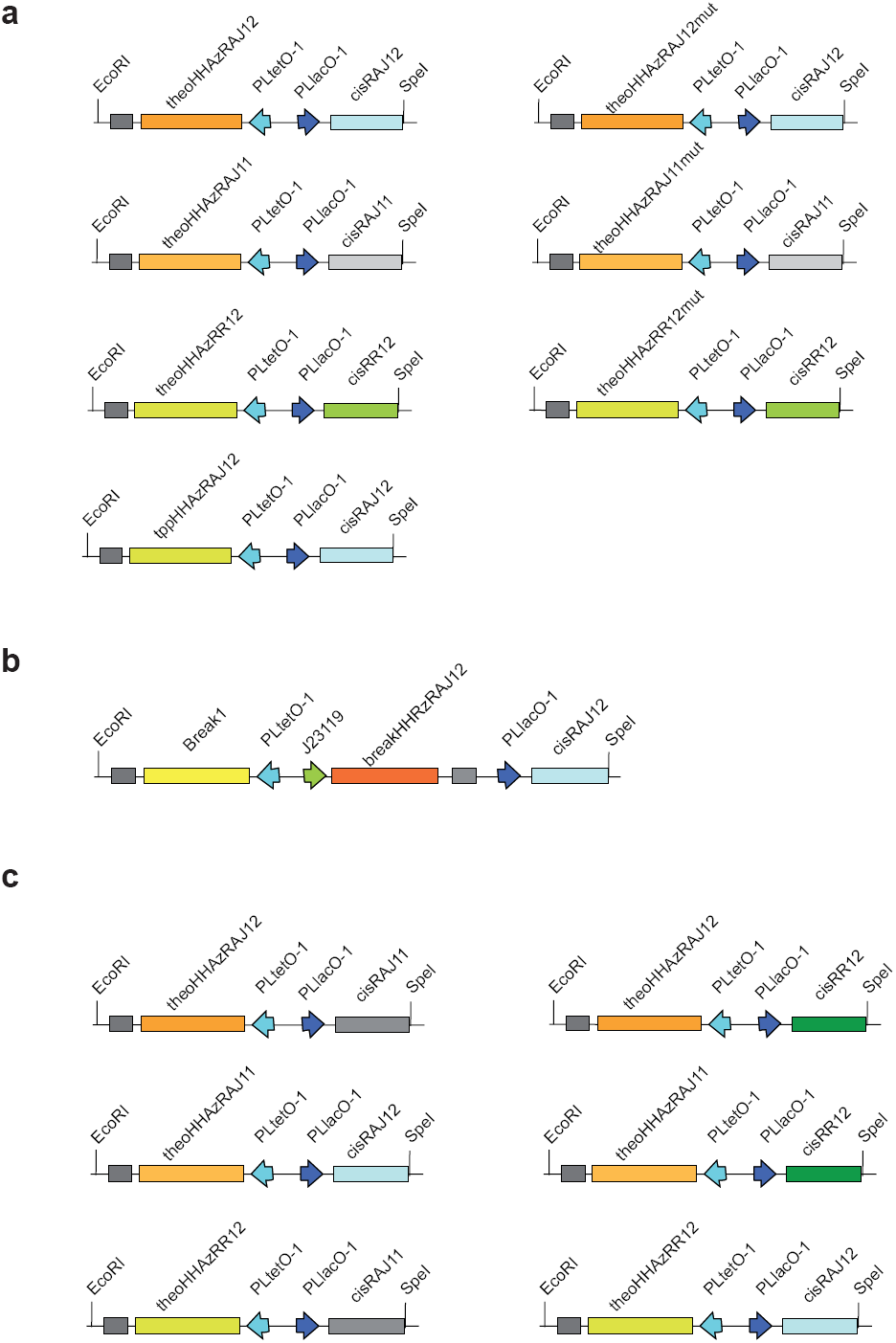
Summary of the gene cassettes used in this work. (**a**) Four small-molecule sensing regazymes were synthesized as indicated in the left panel. The right panel shows the corresponding two-nucleotide inactivation mutants generated by site-directed mutagenesis (see **Supplementary Materials and Methods**). (**b**) One sRNA-sensing regazyme was synthesized as indicated, and a control system. Here, the signal sRNA was placed under the control of P_LtetO1_, the regazyme under the control of the constitutive promoter J23119. (**c**) The gene cassettes used in the orthogonality assay were constructed by using restriction enzyme digestion and T4 ligation. We generated six combinations of regazymes and *cis*-regulators. Maps were constructed with the free software Savvy (http://bioinformatics.org/savvy/).

**Supplementary Figure 3:**
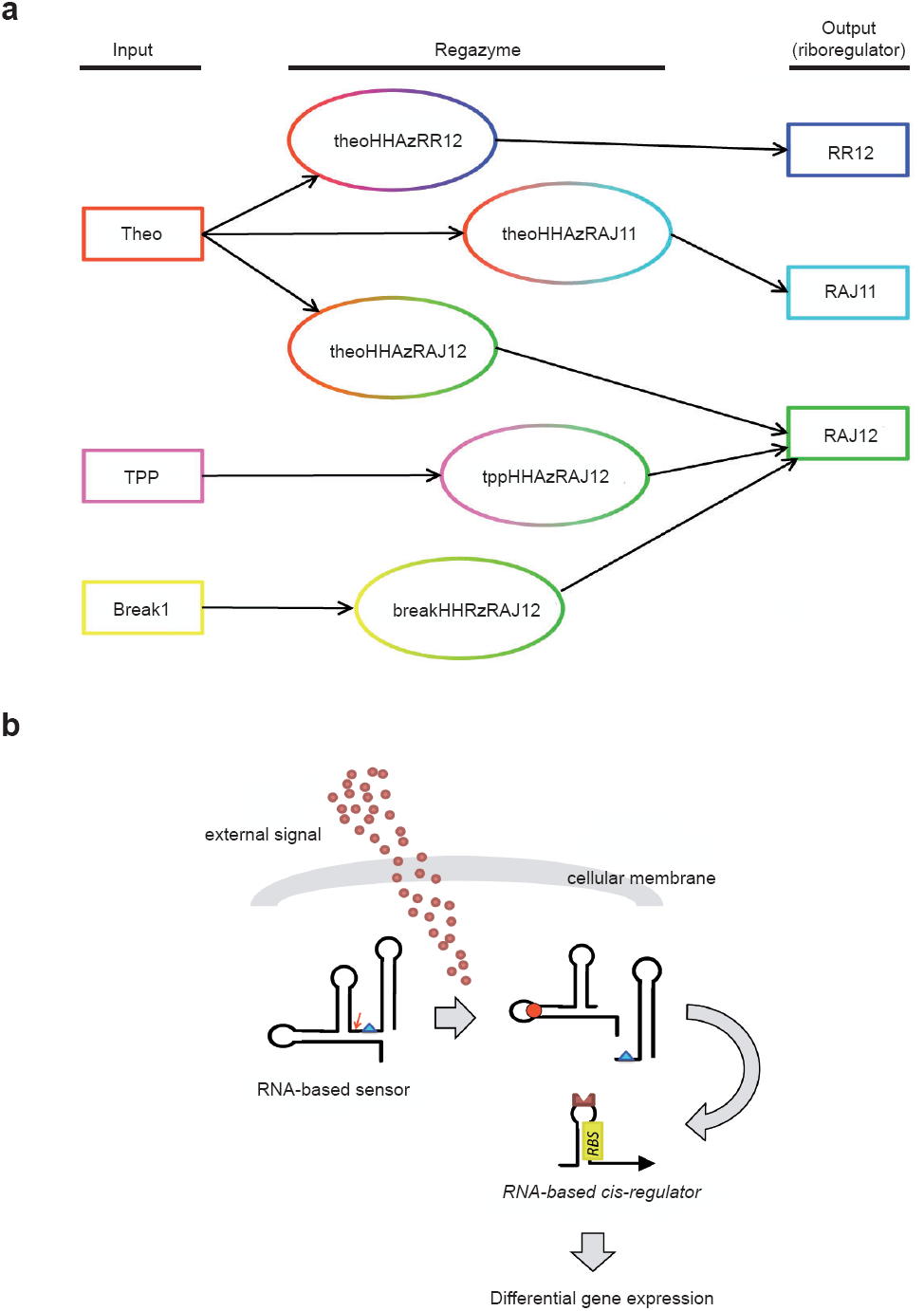
Summary of the signaling pathways generated in this study. (**a**) Regazymes were shown as ellipses with two different colors that correspond to the sensor domain (aptazyme) and the actuator domain (riboregulator). The input signals were two small molecules: theophylline (Theo) and thiamine pyrophosphate (TPP), and one sRNA: Break1. The actuators corresponded to three different riboregulators: sRNAs from systems RR12, RAJ11, and RAJ12, which link downstream to their corresponding *cis*-repressed gene expression platforms. (**b**) Illustration of a regazyme exploited for sensing and transducing environmental signals.

**Supplementary Figure 4:**
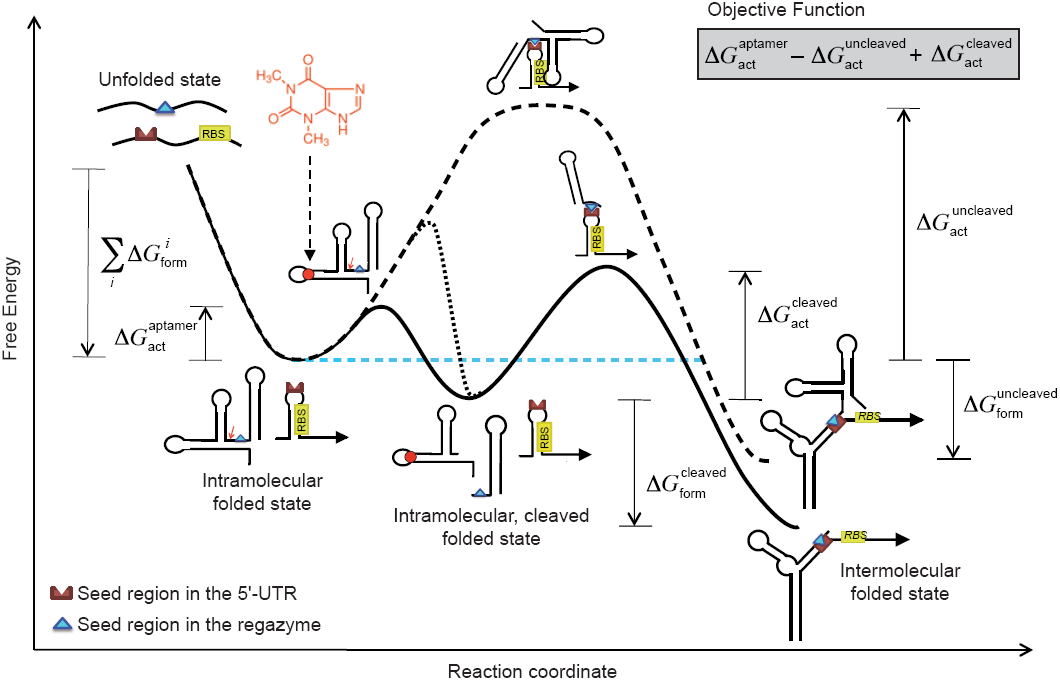
Detailed energy landscape in terms of a reaction coordinate. The different intra- and intermolecular folding states are shown, as well as the cleaved and uncleaved states. All free energies gaps are illustrated. Three different trajectories over the landscape can be followed. One corresponds to the ligand-induced cleavage of the regazyme and subsequent binding of the riboregulator to the mRNA (solid line). Another corresponds to the natural self-cleavage of the regazyme (in absence of ligand) and also subsequent binding of the riboregulator (dotted line).For indication purposes, we present this trajectory in the same landscape, although the associated reaction coordinate would be different. And, finally, another corresponds to the eventual binding of the regazyme (in absence of cleavage) to the mRNA (dashed line).

**Supplementary Figure 5:**
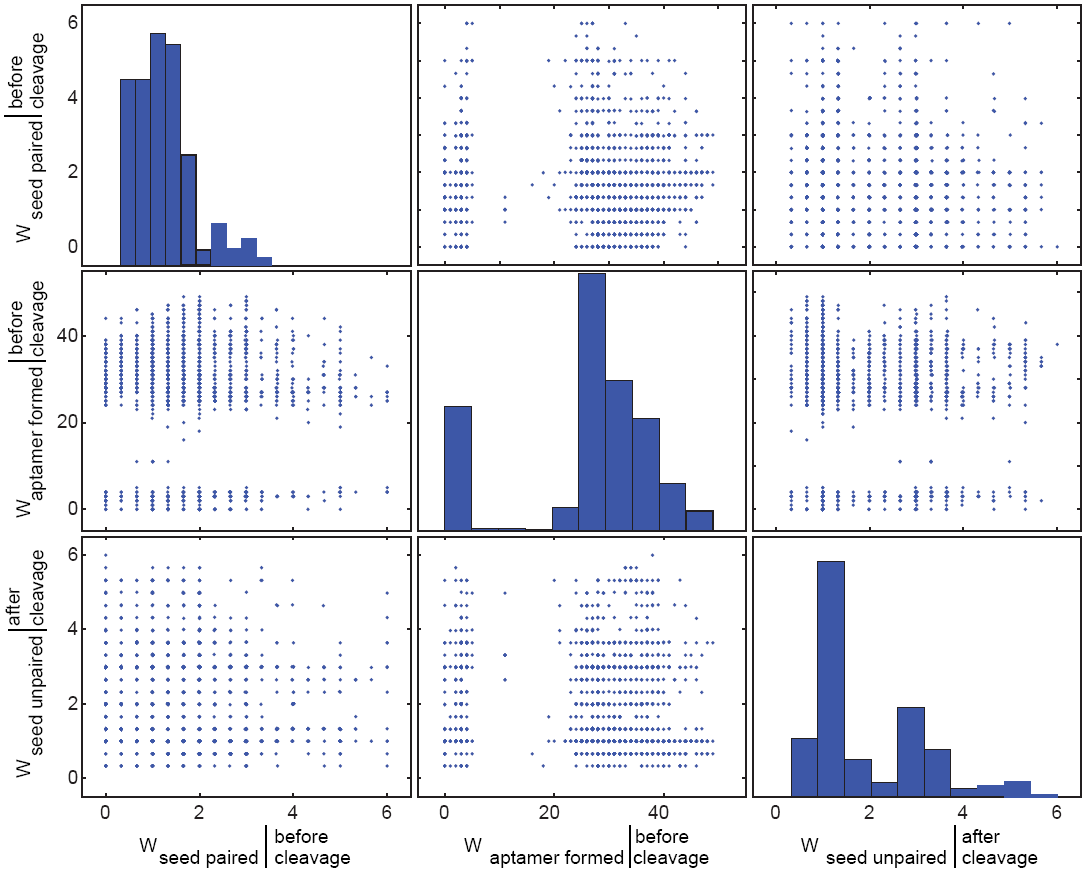
Energy landscape in terms of thermodynamic works (*W*, see Supplementary Materials and Methods). We represent planar projections of main Figure 1d, together with the distributions of the different works (to get the seed paired and the aptamer formed before cleavage, and the seed unpaired after cleavage).

**Supplementary Figure 6:**
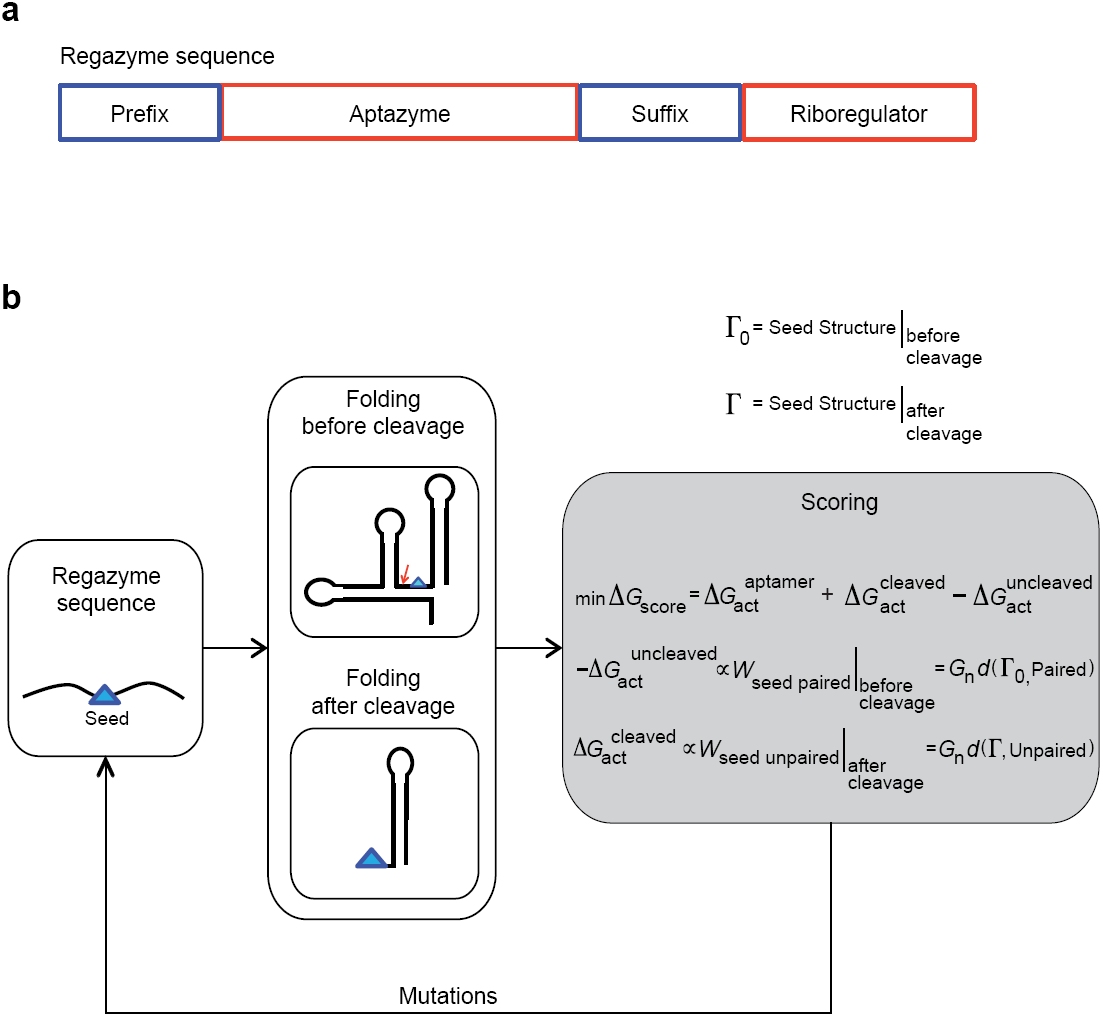
Scheme of the algorithm for regazyme sequence design. (**a**) Modular sequence of a regazyme. It is composed of prefix and suffix sequences flanking an aptazyme plus a riboregulator. The aptazyme and riboregulator sequences are kept fixed. The prefix and suffix are designed to get the intended regulation: in our case, to produce a functional riboregulator after ligand-induced cleavage. (**b**) Optimization scheme for sequence design. Starting from random sequences for the prefix and suffix, the algorithm implements a heuristic optimization (based on Monte Carlo Simulated Annealing) where random mutations (involving replacements, additions, or deletions) are applied and selected with an objective function.

**Supplementary Figure 7:**
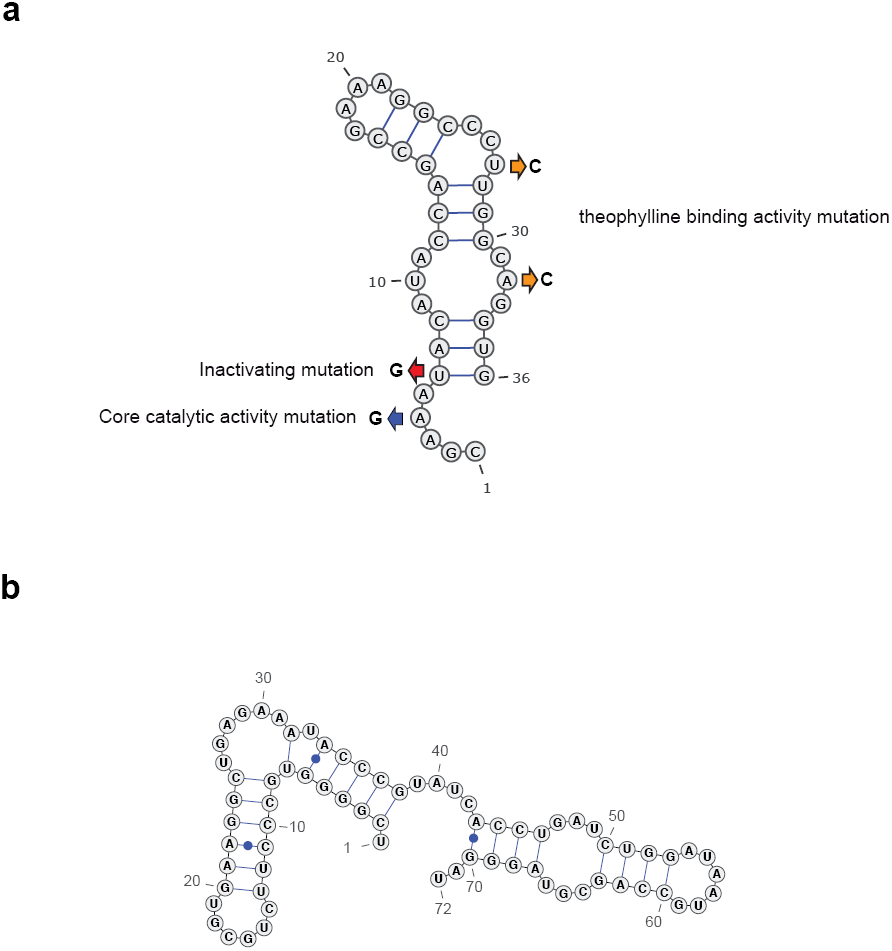
Sequences and secondary structures of the aptamers considered in this work. We designed regazymes with (**a**) theophylline and (**b**) thiamine pyrophosphate aptamers. In addition, we constructed three different types of mutants in the theophylline aptamer, indicated as arrows in (**a**), as a negative control and also to study the natural self-cleavage of the regazyme. This way, we affect the core catalytic activity of ribozyme (blue arrow A to G mutation in **a**), inactivating ribozyme (red arrow U to G mutation in **a**) and binding affinity to the ligand (orange arrow U, A to C mutation in **a**).

**Supplementary Figure 8:**
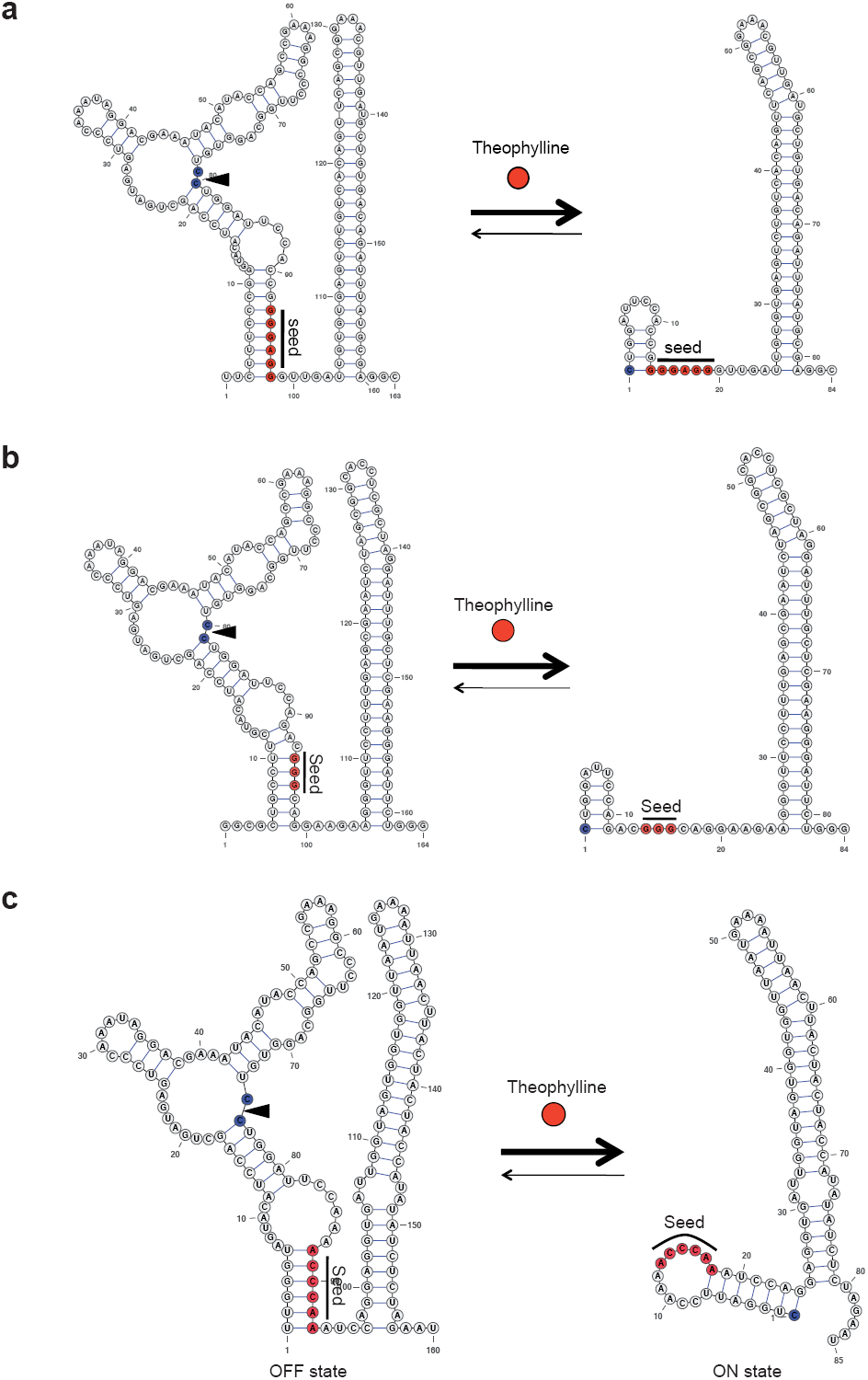
Structural detail of the catalytic reaction of the regazymes. The catalytic site (for self-cleavage) was shown in blue, and marked with an arrow. In the OFF state, the seed (shown in red) is complementary paired. However, in the ON state (after cleavage), it is unpaired and then exposed to the solvent. The secondary structures were predicted by NUPACK (without pseudoknots)^12^ and then plotted with VARNA^13^. (**a**) theoHHAzRAJ11; (**b**) theoHHAzRAJ12; and (**c**) theoHHAzRR12.

**Supplementary Figure 9.**
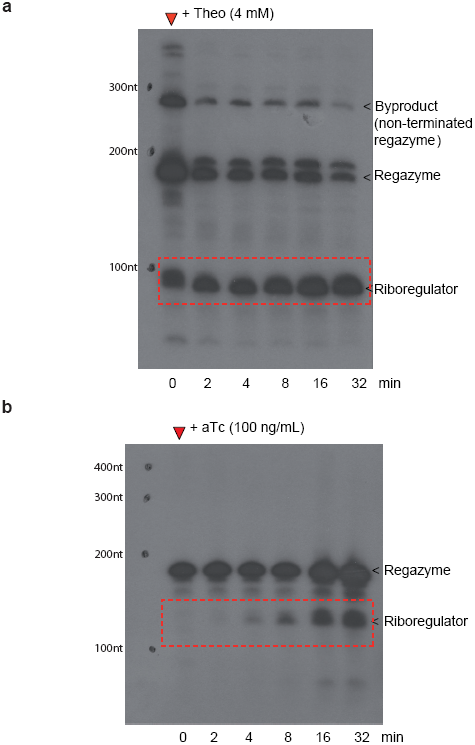
The complete gel images of *in vitro* northern blot corresponding to Figures 2 and 3. RNA extracts from bacterial cells treated with ligands for indicated time points were processed with [^32^P]-labelled RNA probes that recognize regazyme sequences, and separated on northern blot for quantification of the cleavage detailed in **Supplementary Materials and Methods**. (**a**) theoHHAzRAJ12; (**b**) breakHHRzRAJ12. The strong secondary structure of the RNAs provokes a slight displacement with respect to the markers. Note that the probe is the same for the two gels and corresponds to the reverse complement of regazyme theoHHAzRAJ12. In (**a**), comparison with *in vitro* results suggests that the 5’ fragment is quickly degraded *in vivo*. In (**b**), only one product is detected (riboregulator) as the probe cannot reveal the 5’ fragment. The products were shown by red dashed box. Note that these gels have been repeated three times each, obtaining always the same results. On the one hand, regazyme theoHHAzRAJ12 has a length of 194 nt. The band corresponding to the riboregulator released after cleavage, of 114 nt, appears to migrate faster. On the other hand, regazyme breakHHRzRAJ12 has a length of 196 nt. The band corresponding to the riboregulator released after cleavage, of 112 nt, migrates a bit slower. Our results suggest nevertheless that these bands correspond to products of the cleavage reactions in the presence of the ligands (no bands are seen in absence of them).

**Supplementary Figure 10.**
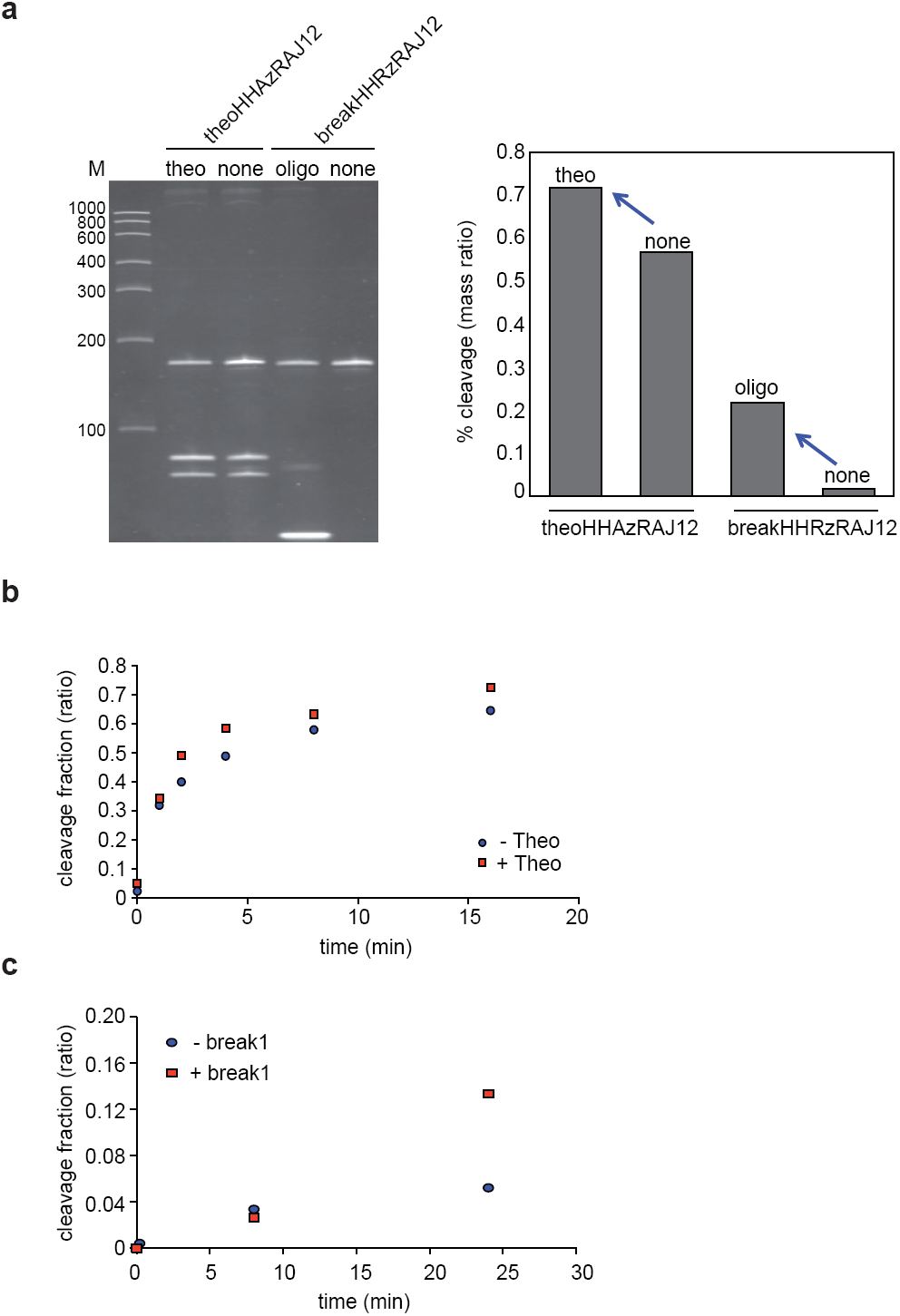
*In vitro* transcribed regazyme self-cleavage assay. (**a**) Regazymes theoHHAzRAJ12 and breakHHRzRAJ12 were *in vitro* transcribed with or without corresponding ligands (theo: 4 mM theophylline; oligo: 3 μM Break1 oligo). The products were separated on PAGE gels followed by ethidium bromide. For theoHHAzRAJ12 two bands (85nt and 92nt) were expected as cleaved products, however, for breakHHRzRAJ12 two bands (89nt and 90nt) were expected as cleaved products. Quantification was shown in the right panel based on the mass ratio between cleaved products and precursors. M: RNA ladders. (**b,c**) *In vitro* time-course of self-cleavage activity assay for regazyme theoHHAzRAJ12 (**b**) and breakHHRzRAJ12 (**c**). The self-cleavage activity was monitored with denaturing PAGE in the presence or absence of ligands at indicated time points. The quantification of cleavage ratio (cleaved / total) is shown here. In the *in vitro* assay, RNA is not produced nor degraded during the cleavage, whereas *in vivo* RNA is produced, degraded and cleaved at the same time. Therefore, the expected kinetics are different. The cleaved fraction increases with time upon the addition of the ligand. Without the presence of ligand, the regazyme is still cleaved *in vitro* (leakage activity), but *in vivo* system is in a steady state. This is also shown by our mathematical model for the dynamics of the system species *in vivo*.

**Supplementary Figure 11:**
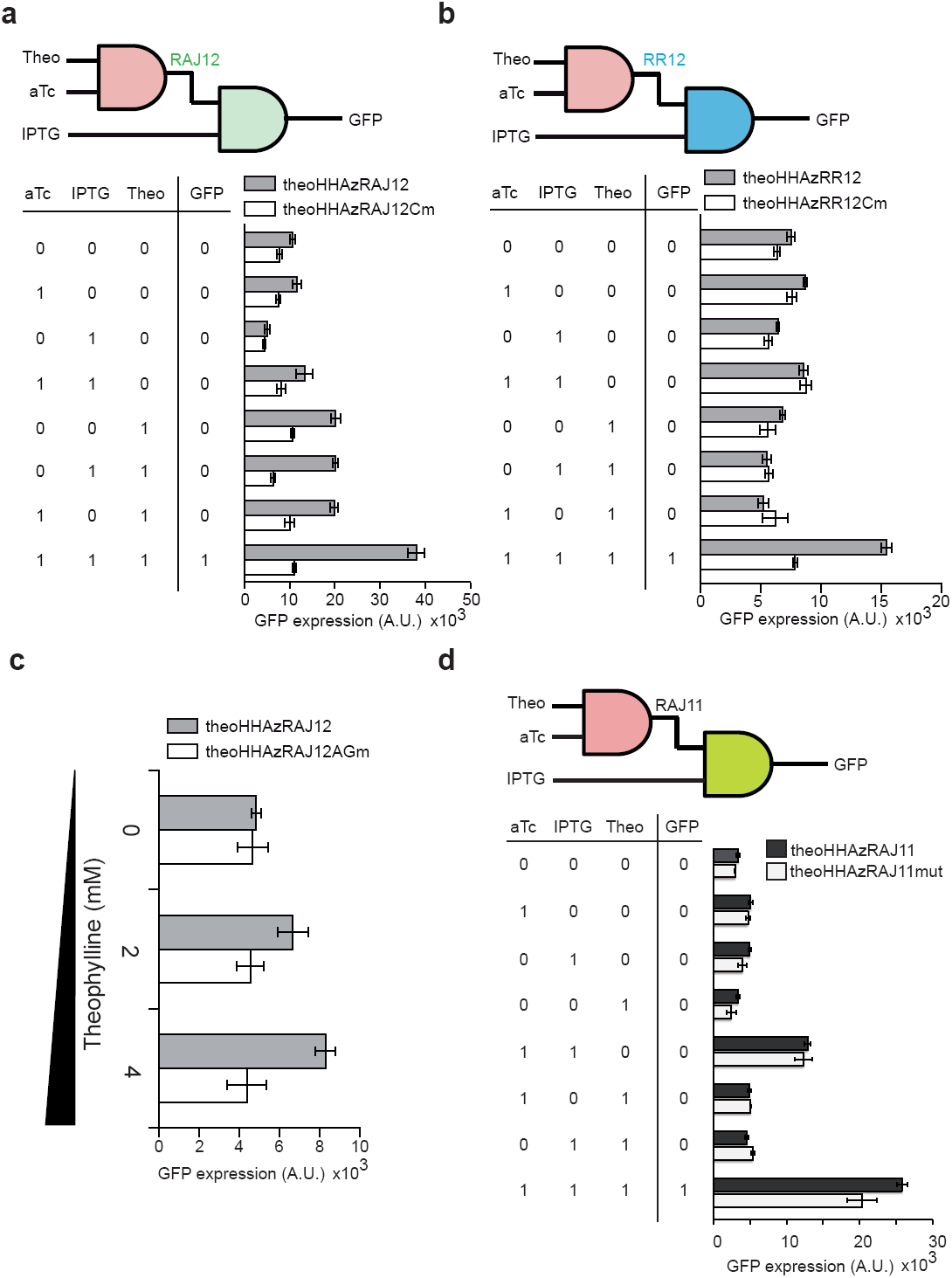
Digital diagram of the regazyme theoHHAzRAJ12, theoHHAzRR12 and theoHHAzRAJ11 together with their characterizations. We present the results of GFP expression (population level) of MG1655Z1 cells expressing Regazyme or Regazyme mutation as control. These were characterized under different combinations of inducers. The truth table is shown in agreement to the experimental data. (**a**) theoHHAzRAJ12 and theoHHAzRAJ12Cm (corresponding to inactivating mutation as shown in Supplementary Figure 7a). (**b**) theoHHAzRR12 and theoHHAzRR12Cm. (**c**) Core catalytic mutation of theoHHAzRAJ12 (theoHHAzRAJ12AGm, see also Supplementary Figure 7a) was analyzed in JS006 strain with different concentrations of theophylline. (**d**) theoHHAzRAJ11 and theoHHAzRAJ11mut (corresponding to theophylline binding activity mutation as shown in Supplementary Figure 7a). The concentrations of inducers were: aTc (100 ng/mL), IPTG (1 mM), and Theo (4 mM). Error bars were standard deviations of three replicates.

**Supplementary Figure 12:**
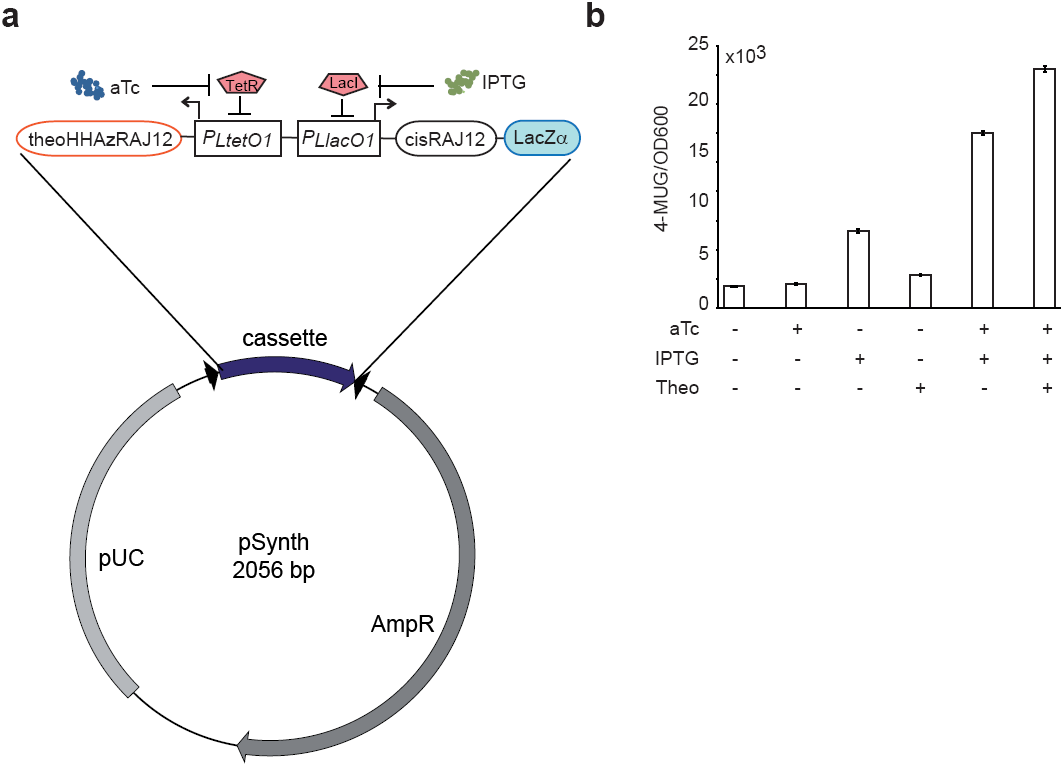
Characterization of regazyme theoHHAzRAJ12 with LacZα reporter. (**a**) The regazyme and reporter gene were synthesized in the commercial plasmid vector pSynth. DH5αZ1 strain was transformed with the plasmid. (**b**) Experimental results. The expression of the reporter gene was characterized thanks to the LacZ enzymatic activity by using the 4-Methylumbelliferyl-β-D-galactopyranoside (4-MUG, Sigma) fluorescence assay^14^. aTc: 100 ng/mL, IPTG: 1 mM, Theo: 4 mM. Error bars were standard deviations of three replicates.

**Supplementary Figure 13:**
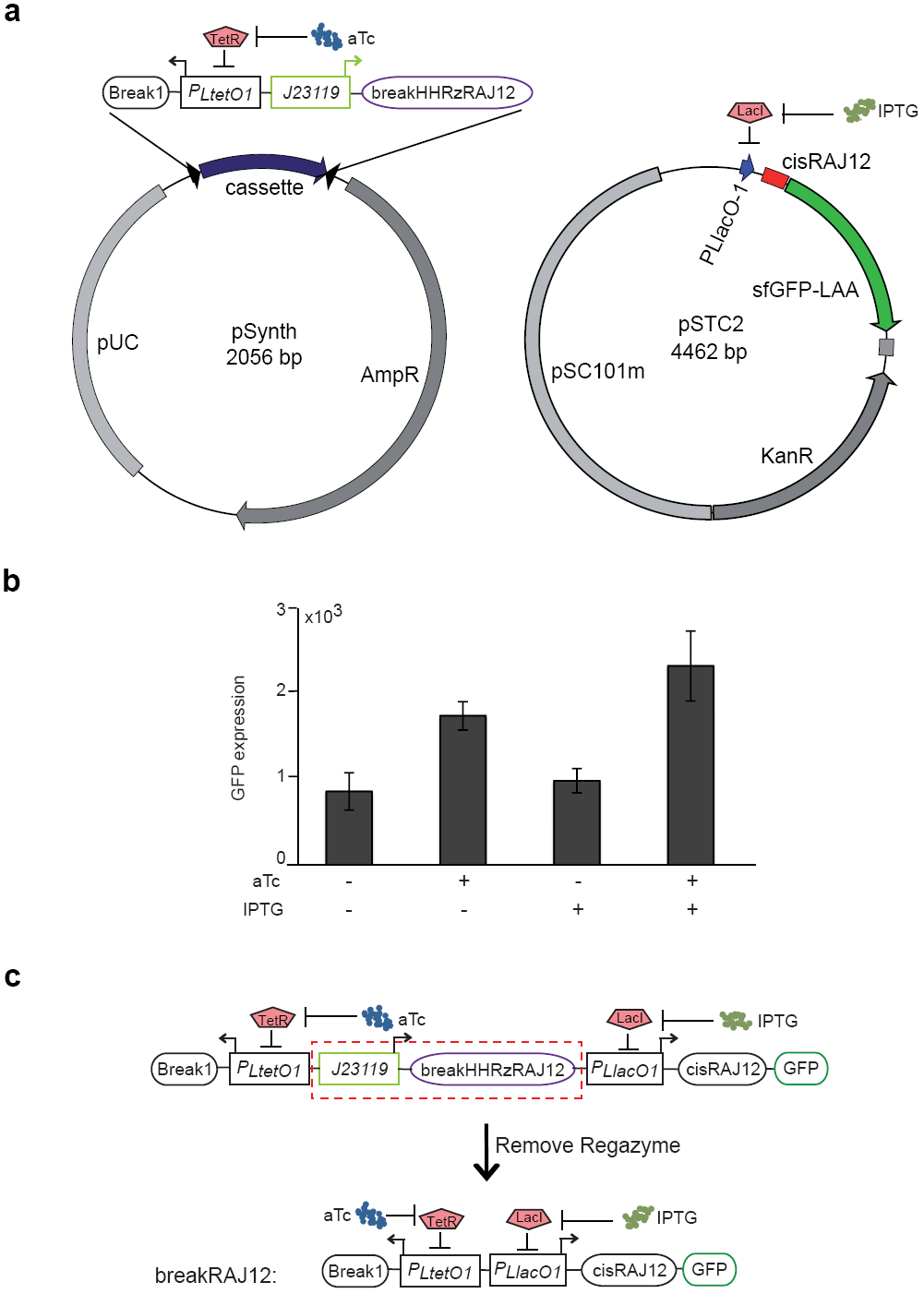
Scheme of the sRNA-sensing regazyme breakHHRzRAJ12 and its dysfunctional mutant. (**a**) The regazyme was synthesized in the commercial plasmid vector pSynth, and it was directly co-transformed with the pSTC2 plasmid expressing the *cis*-repressed sfGFP reporter in MG1655Z1 cells. (**b**) The gene expression was characterized through using the reporter sfGFP fluorescence for cells co-transformed with the two plasmids. We observe lower activity than for a system expressed from a single plasmid which is consistent to reduced interaction ability between two RNAs expressed from two different plasmids. aTc: 100 ng/mL; IPTG: 1 mM. (**c**) To exclude the possibility that the sRNA *Break1* could directly activate the *cis*-repressed sfGFP, we generated a dysfunctional mutant by removing the regazyme (breakHHRzRAJ12) from the plasmid by PCR-based mutagenesis. The corresponding characterization at population level was shown in Figure 3b.

**Supplementary Figure 14:**
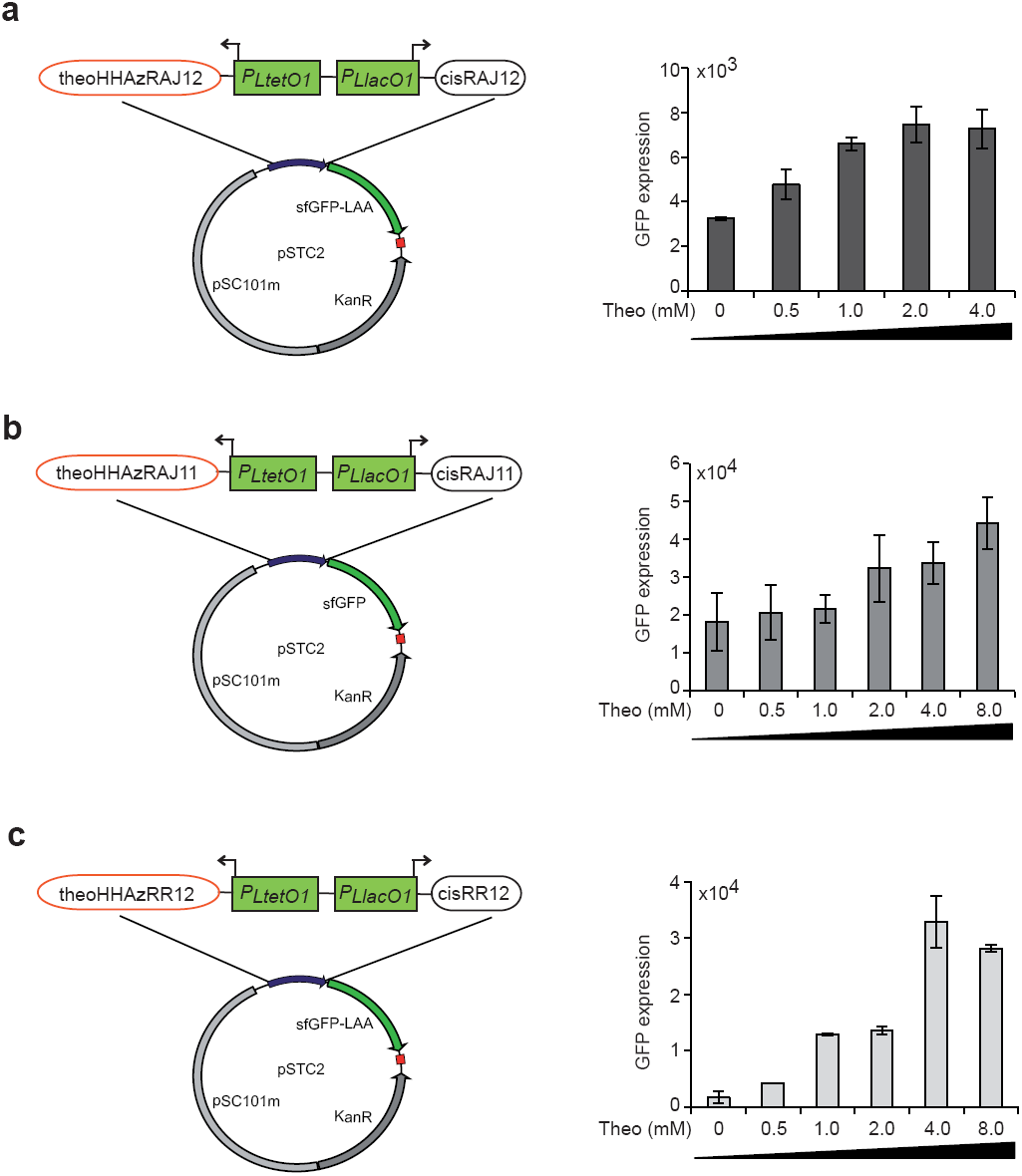
Dose-dependent activation of the three theophylline sensing regazymes. The left panel shows the construct that has been used for the characterization, while the right panel shows the GFP expression in MG1655Z1 cells expressing the corresponding plasmid. (**a**) theoHHAzRAJ12; (**b**) theoHHAzRAJ11; (**c**) theoHHAzRR12. Theo: theophylline. Error bars were standard deviations of six replicates.

**Supplementary Figure 15:**
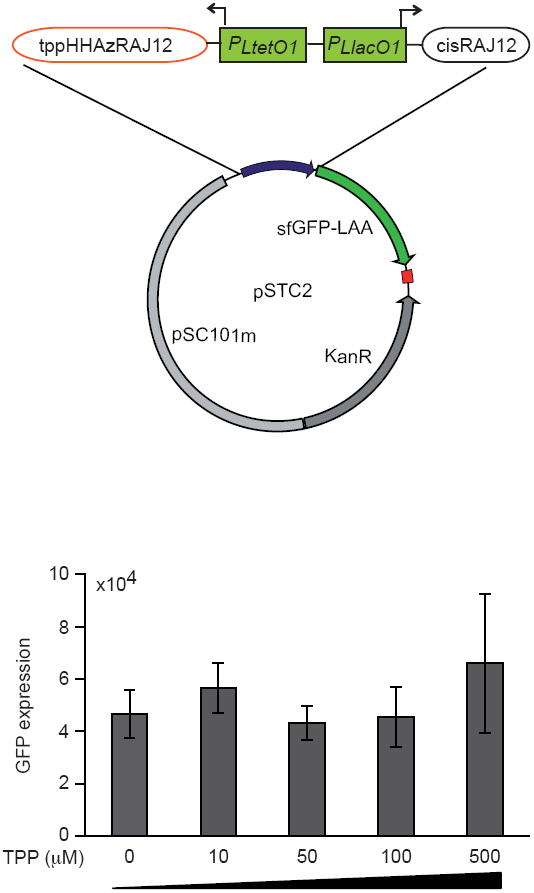
Characterization of regazyme tppHHAzRAJ12. We present the results of GFP expression (population level) of T7 express cells (New England bioLAB, USA) expressing tppHHAzRAJ12. The construction being used for the characterization was shown in the upper panel. T7 express cells (New England BioLAB, USA) transformed with the construct were exposed to different concentration of thiamine as indicated in the figure. Error bars were standard deviation of three replicates.

**Supplementary Figure 16:**
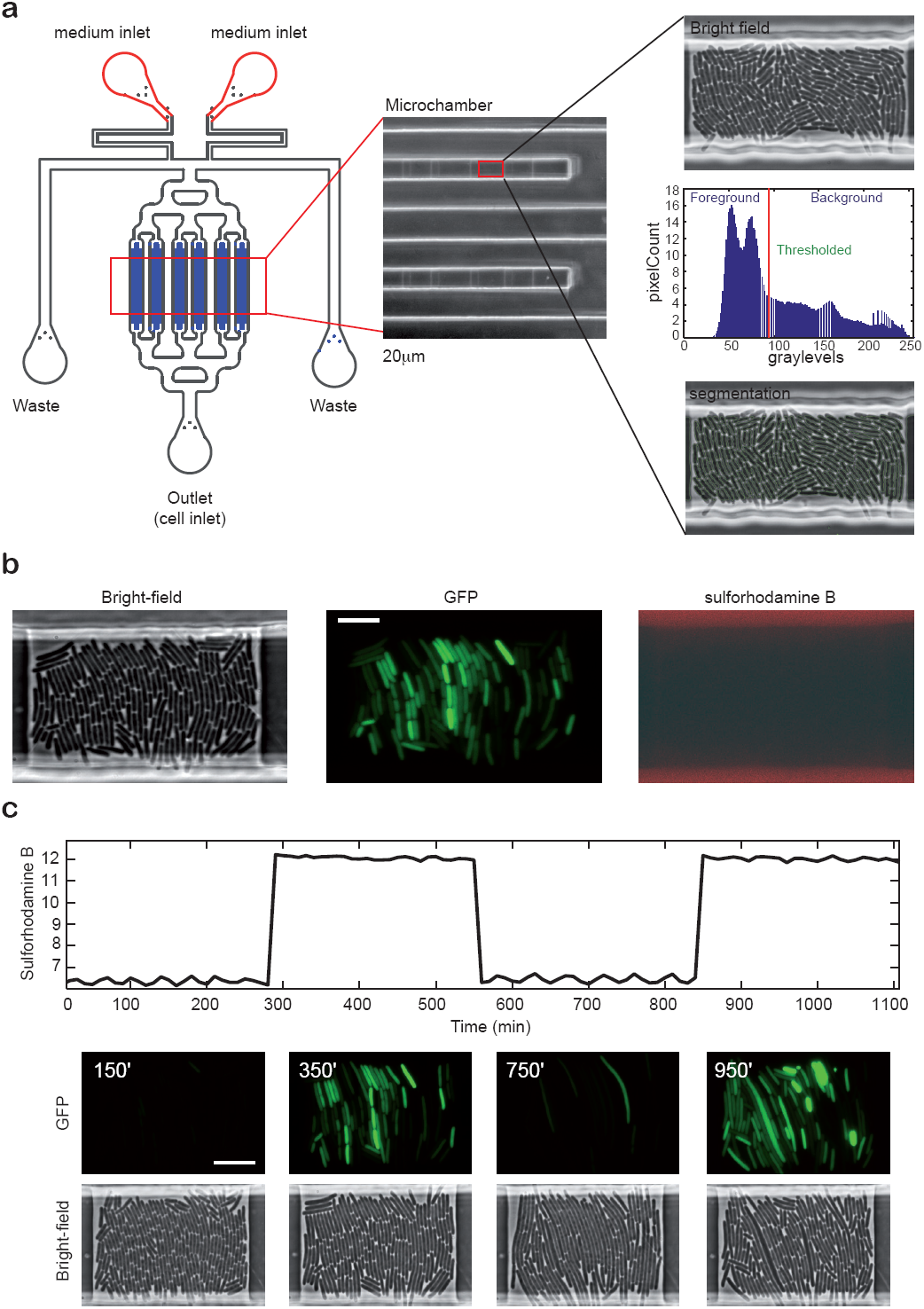
Microfluidics device constructed for this study. (**a**) Scheme of the device used for the single cell dynamic analysis in the left panel. The blue part is the microchamber region. An exemplifying image of the microchamber is shown in the middle panel. The microchamber is about 30 μm × 30 μm width and 1 μm height. Cells are loaded from the cell inlet and trapped in the microchamber. Exemplifying images of cell trapping are shown in the right panel. Images are then analyzed (thresholding and segmentation, see **Supplementary Materials and Methods**). (**b**) Cell images from the bright-field and fluorescence (green and red) channels. Bright-field images can serve for segmentation and tracking. In our experiments, the medium containing the inducer (e.g., theophylline) also contained the red fluorescent dye sulforhodamine B. The dynamics of red fluorescence was therefore the same as the one of the inducer, which serves to account for the diffusion of the molecules in the microchamber. (**c**) Exemplifying images to demonstrate that GFP expression correlates well with the inducer amount into the microchambers. On the top, we show a time-dependent measurement of the red fluorescence intensity in the microchamber. On the bottom, we show cell images of green fluorescence and bright field in the same time scale. Bar: 10 μm.

**Supplementary Figure 17:**
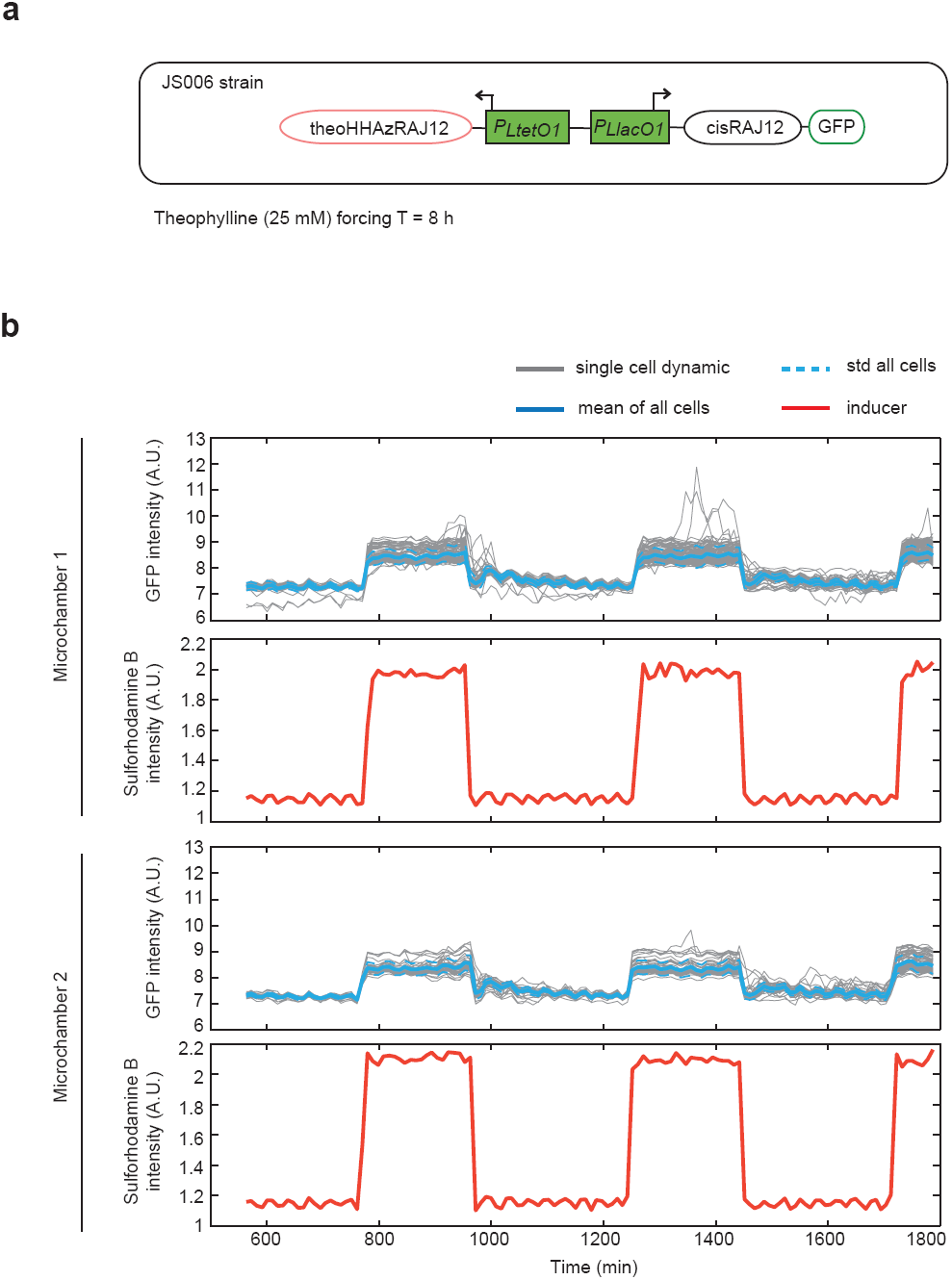
Microfluidics-based single cell analysis of the theophylline-sensing regazyme theoHHAzRAJ12. (**a**) Scheme of the genetic circuit in JS006 cells. Although the regazyme and mRNA were under the control of inducible promoters (P_LtetO1_ and P_LlacO1_ respectively), in JS006 strain there is no expression of repressors LacI and TetR. Therefore, cells could be induced with theophylline directly. A square wave of theophylline (25 mM) with period T = 8 h (i.e., 4 h induction and 4 h relaxation) was applied. (**b**) Single cell tracking in two different and independent microchambers. Data and plots were generated with MATLAB (MathWorks).

**Supplementary Figure 18:**
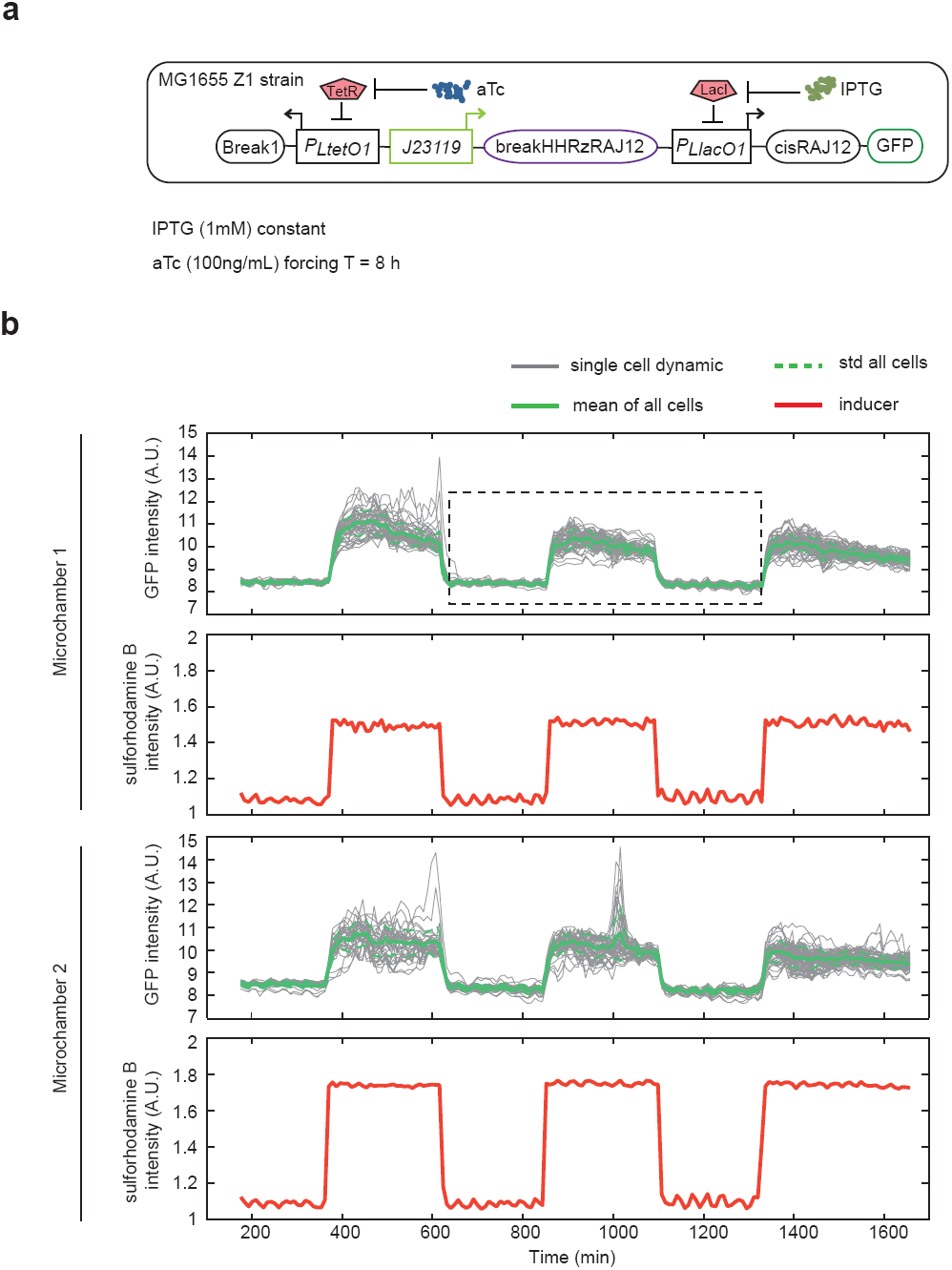
Microfluidics-based single cell analysis of the sRNA-sensing regazyme breakHHRzRAJ12. (**a**) Scheme of the genetic circuit in MG1655Z1 cells. A constant amount of IPTG (1 mM) was established during the whole experiment. A square wave of aTc (100 ng/mL) with period T = 8 h (i.e., 4 h induction and 4 h relaxation) was applied. (**b**) Single cell tracking in two different and independent microchambers. Data and plots were generated with MATLAB (MathWorks).

**Supplementary Figure 19:**
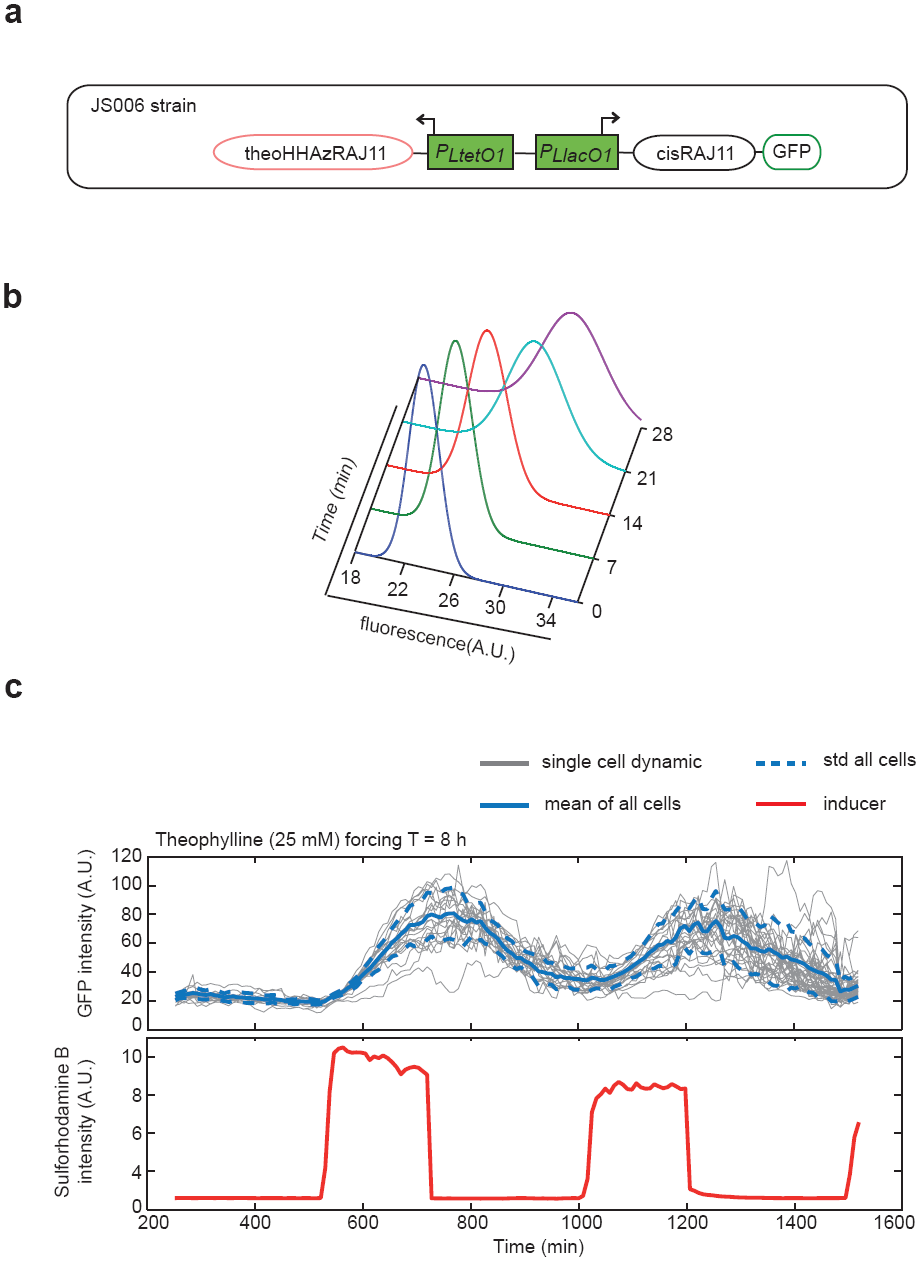
Microfluidics-based single cell analysis of the regazyme theoHHAzRAJ11. (**a**) Scheme of the genetic circuit. (**b**) Distributions of fluorescence with time across a population of cells and fitted with a Gaussian model. (**c**) Single cell tracking as done as in Supplementary Figure 17. Data and plots were generated with MATLAB (MathWorks, USA). JS006 cells were transformed with the corresponding plasmid and were characterized.in the microfluidics device.

**Supplementary Figure 20:**
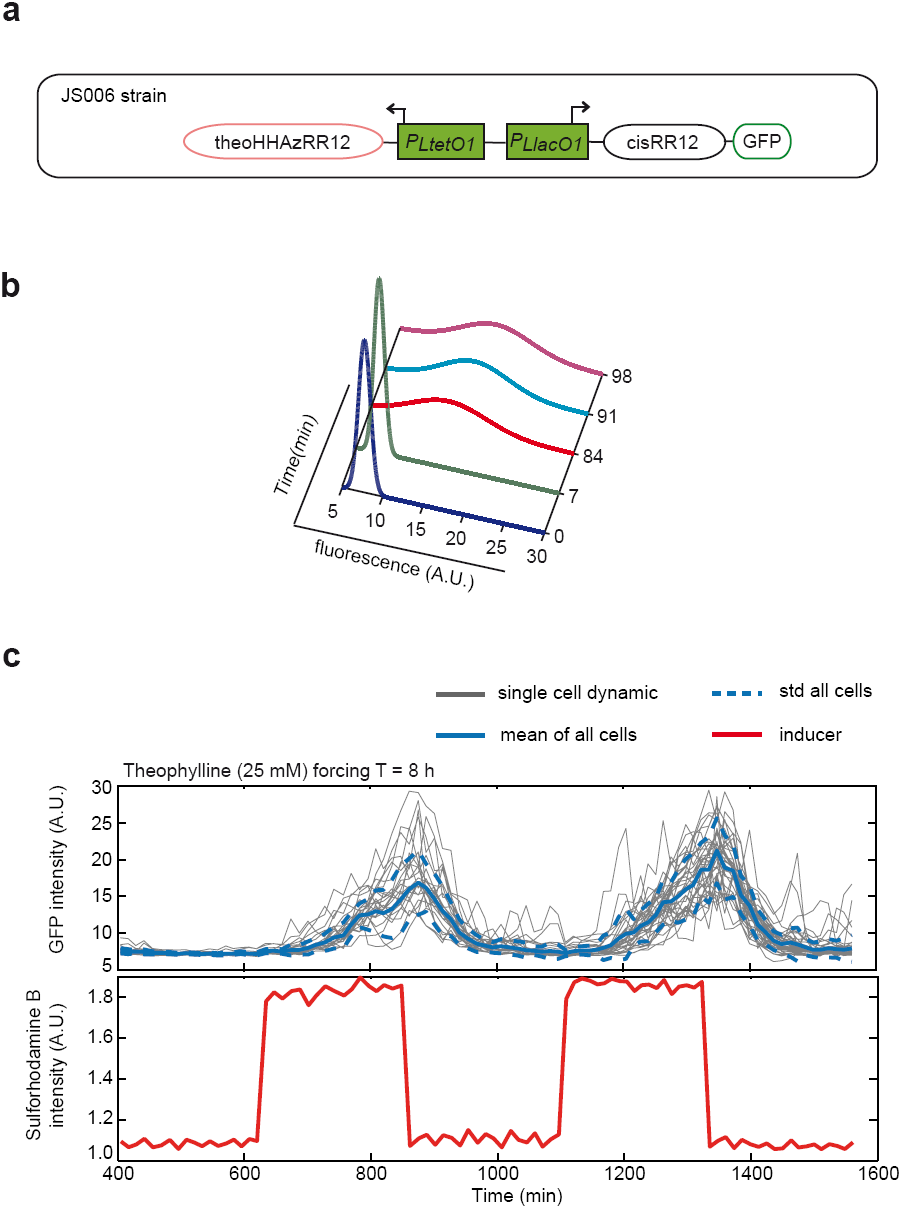
Microfluidics-based single cell analysis of the regazyme theoHHAzRR12. (**a**) Scheme of the genetic circuit. (**b**) Distributions of fluorescence with time across a population of cells and fitted with a Gaussian model. (**c**) Single cell tracking as done in Supplementary Figure 17. Data and plots were generated with MATLAB (MathWorks, USA). JS006 cells were transformed with the corresponding plasmid and were characterized in the microfluidics device.

**Supplementary Figure 21:**
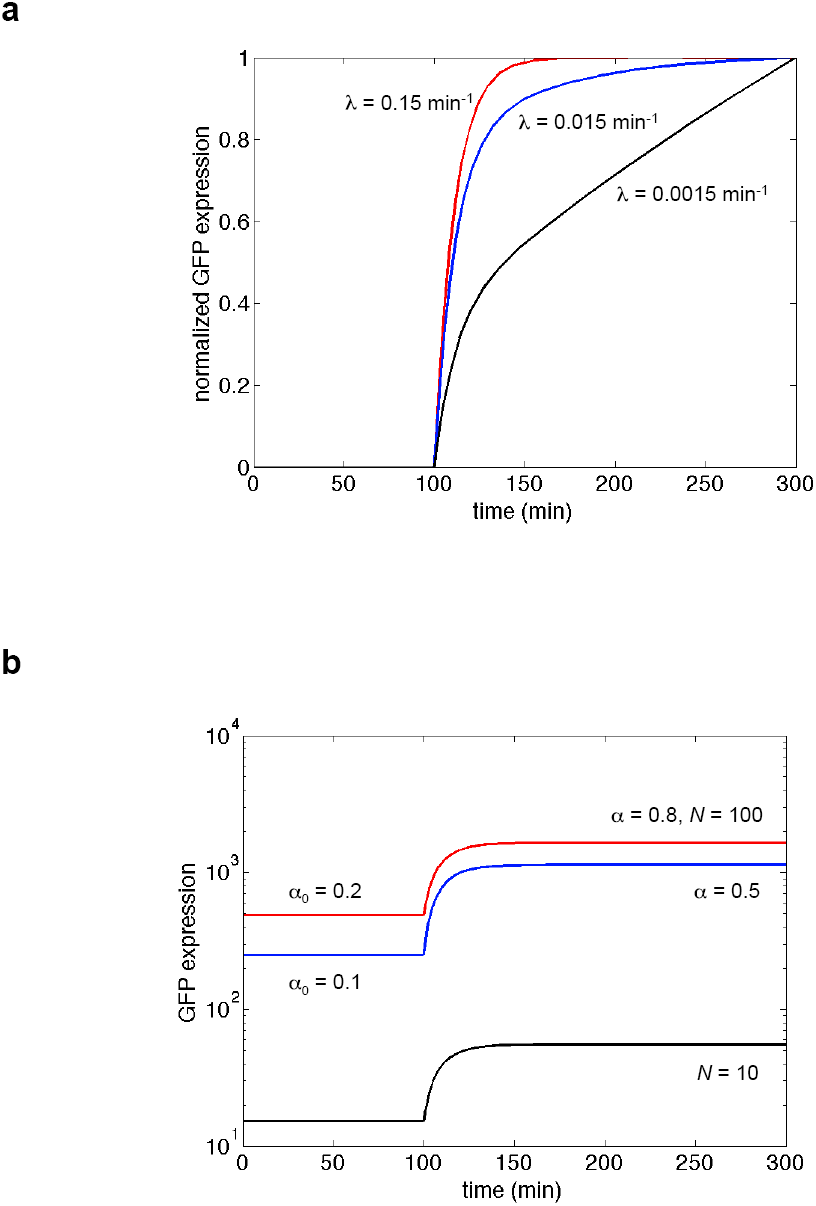
Model simulations for different parameter values. (**a**) We show how the dynamic response of the system changes with the self-cleavage rate of the regazyme (λ). (**b**) We show that the dynamic response is not affected by the fraction of regazyme cleaved (*α*) or the copy number (*N*). These parameters (in addition to *K*_reg_) only affect the level of the steady state. These dynamics collapse all in a single one when using a normalized variable as in panel (**a**). Theophylline is introduced at time 100 min. If not specified, the parameters take the values shown in Supplementary Table 5.

**Supplementary Figure 22.**
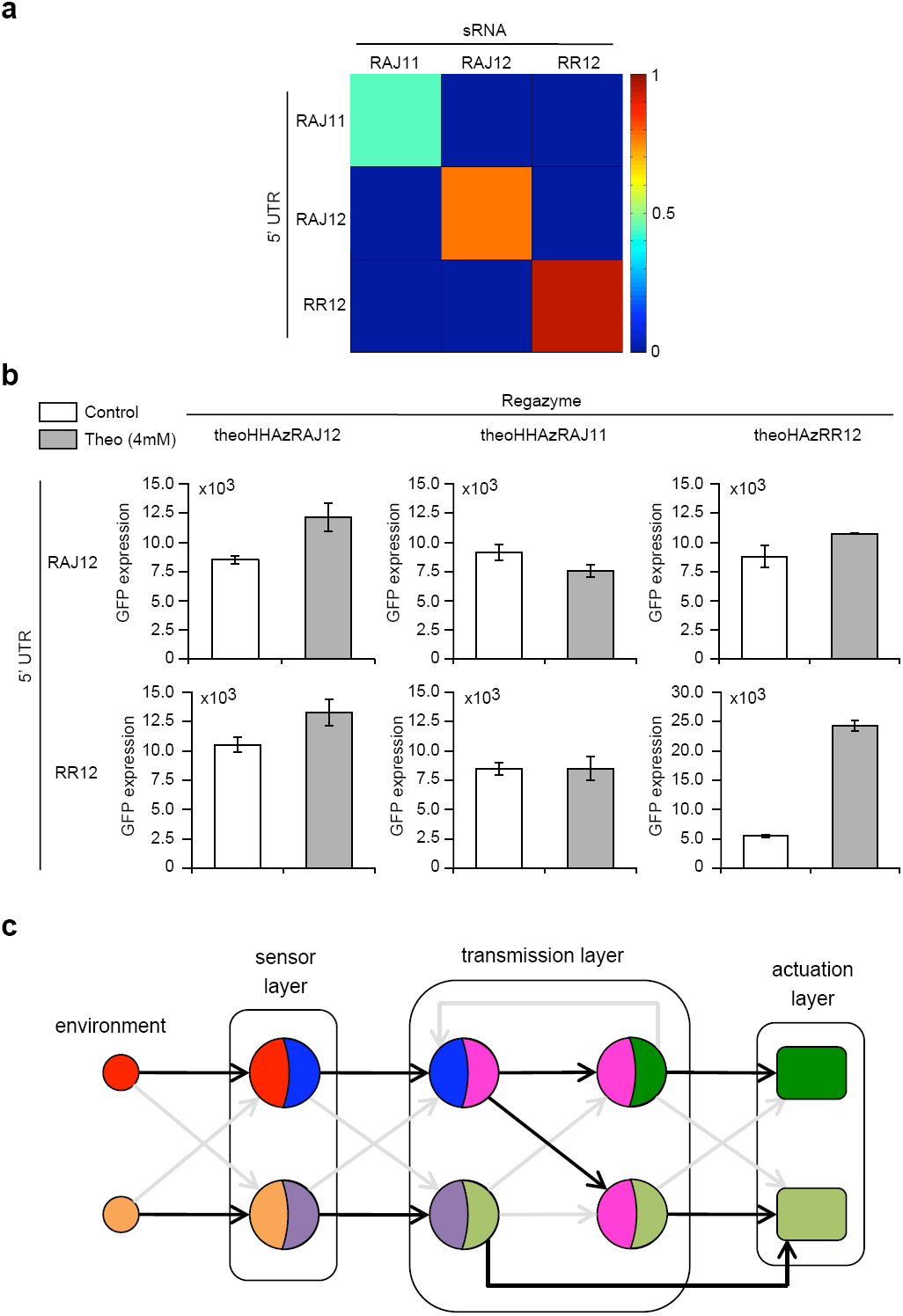
Orthogonality analysis between regazymes. (**a**) Computational prediction of interaction between the released riboregulator and the 5’ UTR (for cognate and non-cognate pairs) ^1^. (**b**) Experimental orthogonal analysis between regazymes. JS006 cells were transformed with corresponding constructs for characeterization of the cross-talk between regazymes. Theo: 4mM. Error bars represent standard deviations of three replicates. (**c**) Scheme of the composability of reazymes to implement several circuits.

**Supplementary Figure 23:**
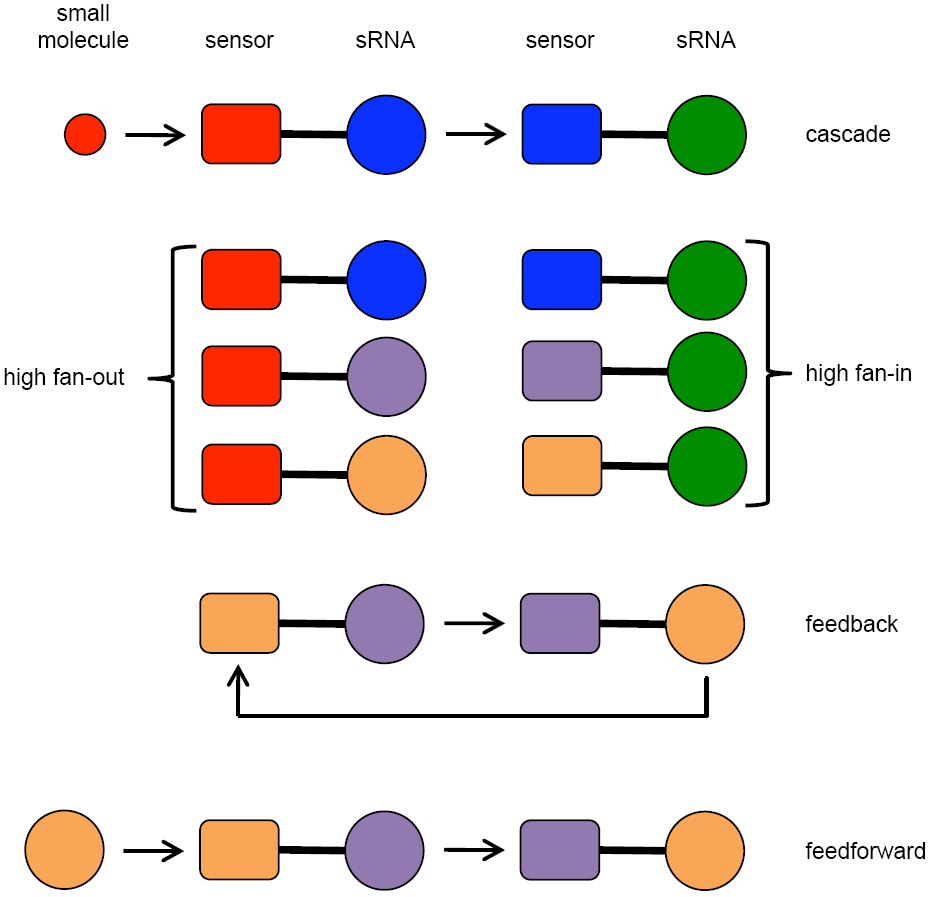
Schemes of expanded application of regazyme-based circuits. We show the implementation of cascades (of small molecule-sensing regazymes coupled with sRNA-sensing regazymes), how to increase the fan-in or fan-out of a regazyme-based system, and also the implementation of feedback and feedforward loops (with sRNA-sensing regazymes).

**Supplementary Tables**

**Supplementary Table 1:**
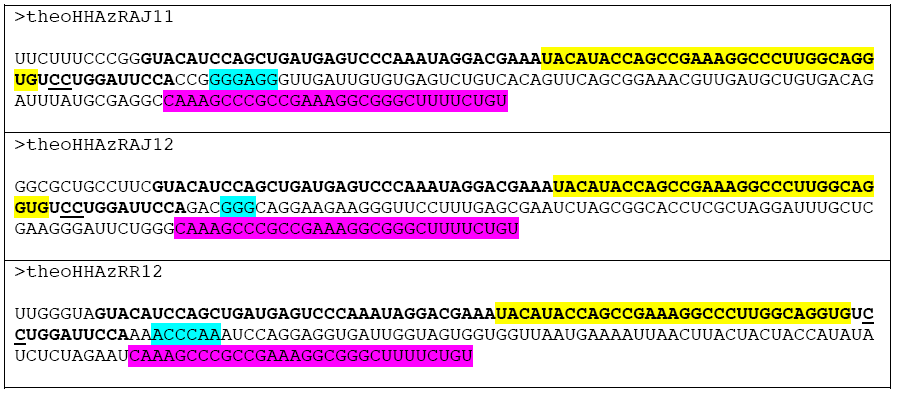
Sequences of regazymes theoHHAzRAJ11, theoHHAzRAJ12 and theoHHAzRR12 designed in this work. Here, theoHHAzX (where X = RAJ11, RAJ12 or RR12) is the chimera between the aptazyme theoHHAz ^15^ and the riboregulator X ^1^,^16^, which can initiate protein translation of an mRNA through activating the appropriate 5’ UTR ^1,16^. The aptazyme sequence is bold-faced with the cleavage site underlined (CC). The theophylline aptamer is shown in yellow. The seed region of the riboregulator is shown in cyan (riboregulators have different seed sequences). The transcription terminator T500 was used in this work ^17^, and it is shown in magenta (efficiency > 90%).

**Supplementary Table 2:**
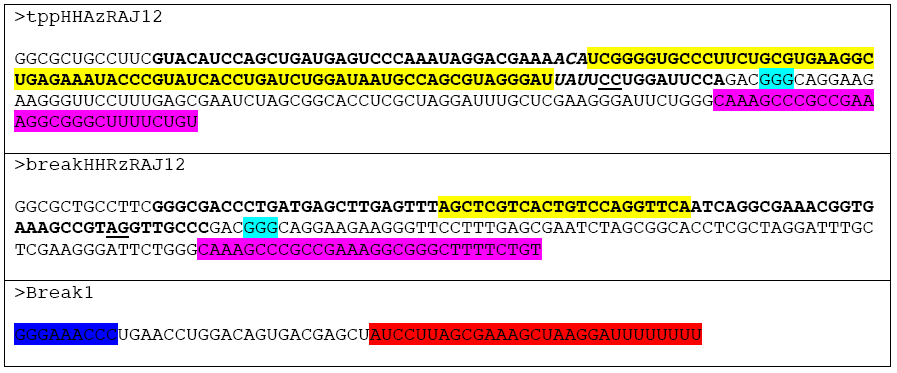
Sequences of regazymes tppHHAzRAJ12 and breakHHRzRAJ12 designed in this work. The tppHHAzRAJ12 is the chimera between the aptazyme tppHHAz ^18^ and the riboregulator RAJ12 ^1^; The breakHHRzRAJ12 is the chimera between a minimal ribozyme ^19^ and the riboregulator RAJ12. The riboregulator can activate protein translation of an mRNA with the appropriate 5’ UTR ^1^. The aptazyme/ribozyme sequence is bold-faced, and the cleavage site is underlined (CC or AG). The thiamine pyrophosphate (TPP) aptamer in tppHHAzRAJ12 and the binding sequence to a small RNA (Break1) in breakHHRzRAJ12 are shown in yellow. The seed region of the riboregulator is shown in cyan. The transcription terminator T500 was used in this work ^17^, and it is shown in magenta (efficiency > 90%).

**Supplementary Table 3:**
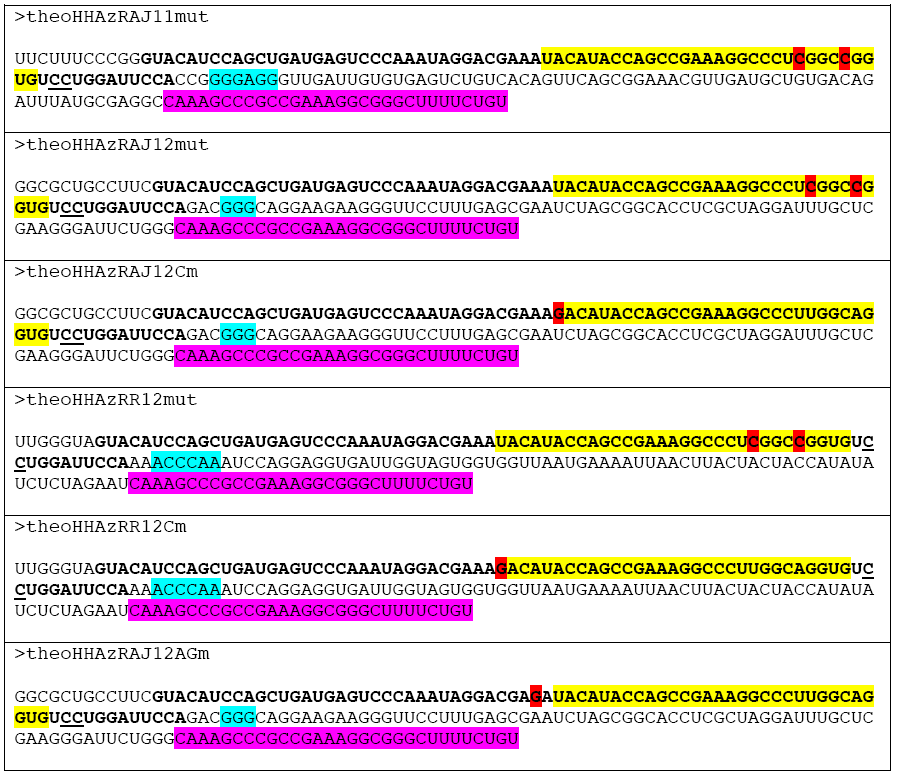
Sequences of dysfunctional mutant regazymes. The theophylline aptamer is shown in yellow (see details in Supplementary Table 1). The mutations U23C and A27C in the aptamer (local numbering) ^20^, which destroy the binding to the ligand, are shown in red. The mutation U to G that inactivates the apatzyme is also shown in red, as well as the mutation A to G in the core catalytic domain destroying the activity.

**Supplementary Table 4:**
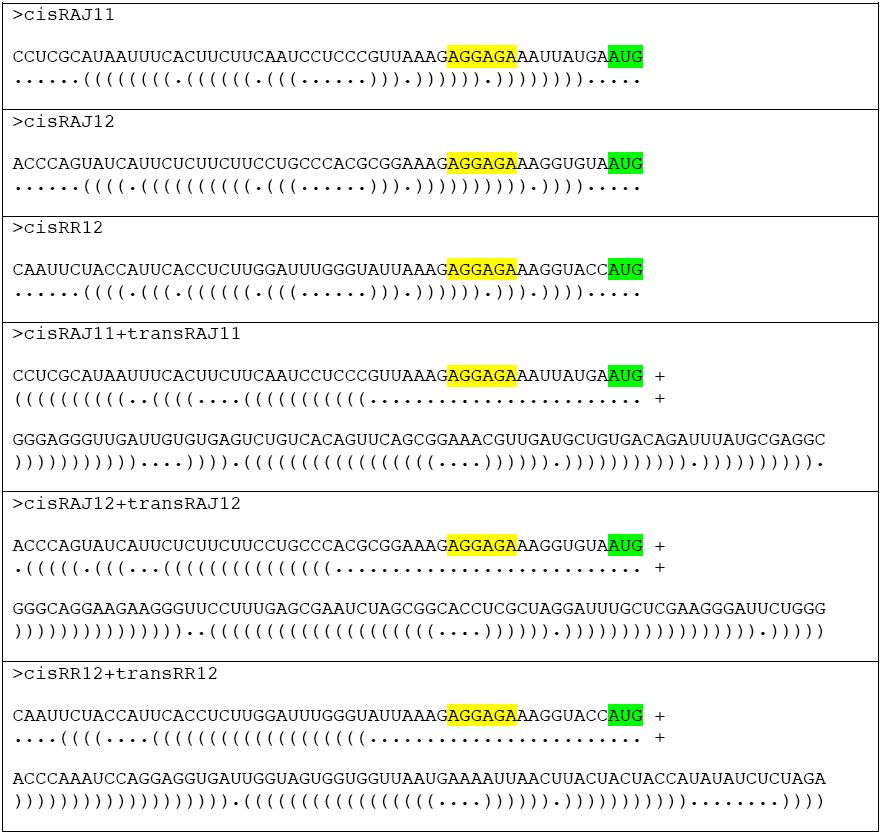
Sequences and structures of the 5’ UTRs. The *cis*-repressed 5’ UTR sequences of the mRNA reporter and the corresponding active complex (5’ UTR + riboregulator) are shown with RBS (Shine-Dalgarno sequence) marked as yellow. The corresponding secondary structures are shown, where RBS is exposed for ribosome binding upon interaction between riboregulator and 5’ UTR ^1^,^16^. The start codon is shown in green.

**Supplementary Table 5:**
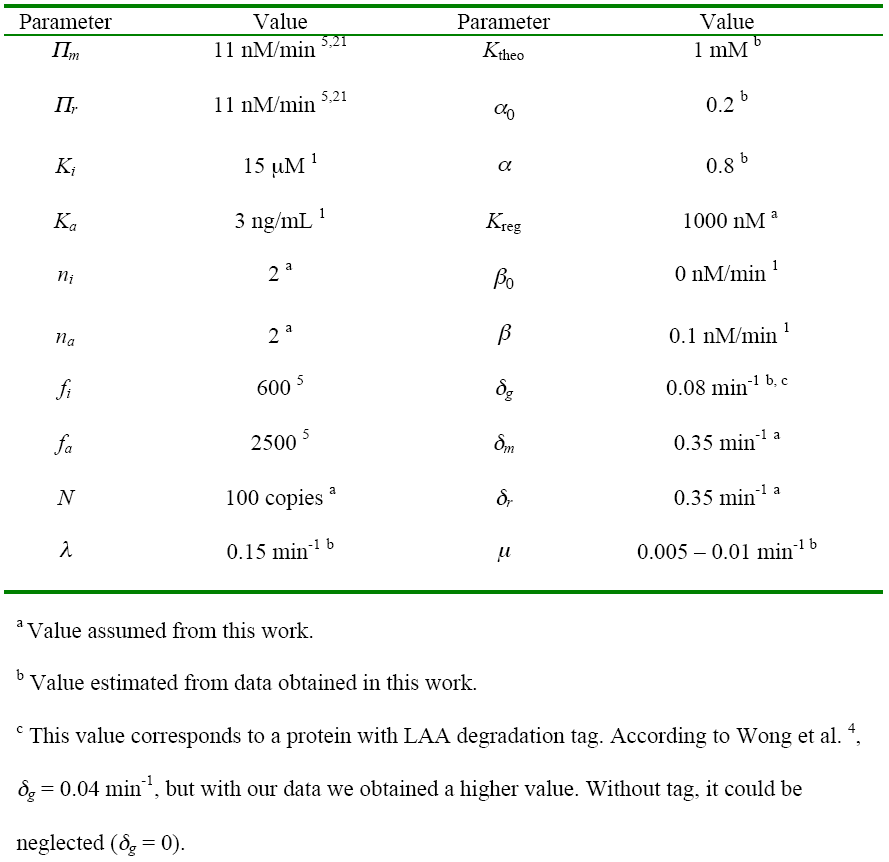
Parameter values for the mathematical model.

**Supplementary Table 6:** Strains and plasmids used in this work.

| Strains or plasmids | Features | Ref. |
| --- | --- | --- |
| *E. coli* DH5α | Commercial | Invitrogen |
| *E. coli* DH5αZ1 | Commercial (DH5α, *lacIQ*, PN25-*tetR*, SpR) | Clontech |
| *E. coli* JS006 | K-12 MG1655, $\Delta araC$, $\Delta lacI$ | Stricker et al <sup>7</sup> |
| *E. coli* MG1655Z1 | K-12 MG1655, *lacIQ*, PN25-*tetR*, SpR | Cox et al <sup>8</sup> |
| *E. coli* T7 Express | Commercial (#C3009I) | NEB |
| pSynth | Commercial plasmid pIDTSMART, pUC ori, AmpR | IDT |
| pSTC1 | pSC101m ori, KanR | Rodrigo et al <sup>1</sup> |
| pSTC2 | pSC101m ori, KanR | This work |
| pSCKtheoRAJ11 | pSTC1, theoHHAzRAJ11, cisRAJ11-sfGFP | This work |
| pSCKtheoRAJ11m | pSTC1, theoHHAzRAJ11mut, cisRAJ11-sfGFP | This work |
| pSCKtheoRAJ12 | pSTC2, theoHHAzRAJ12, cisRAJ12-sfGFP | This work |
| pSCKtheoRAJ12m | pSTC2, theoHHAzRAJ12mut, cisRAJ12-sfGFP | This work |
| pSCKtheoRAJ12Cm | pSTC2, theoHHAzRAJ12Cm, cisRAJ12-sfGFP | This work |
| pSCKtheoRAJ12AGm | pSTC2, theoHHAzRAJ12AGm, cisRAJ12-sfGFP | This work |
| pSCKtheoRR12 | pSTC2, theoHHAzRR12, cisRR12-sfGFP | This work |
| pSCKtheoRR12m | pSTC2, theoHHAzRR12mut, cisRR12-sfGFP | This work |
| pSCKtheoRR12Cm | pSTC2, theoHHAzRR12Cm, cisRR12-sfGFP | This work |
| pSCKtppRAJ12 | pSTC2, tppHHAzRR12, cisRAJ12-sfGFP | This work |
| pUAbreak12 | pIDTSMART, breakHHRzRAJ12, Break1 | This work |
| pUAbreak12m | pIDTSMART, Break1 only | This work |
| pSCKbreak12 | pSTC2, cisRAJ12-sfGFP | This work |
| pSCKtheoRAJ11-o12 | pSTC2, theoHHAzRAJ11, cisRAJ12-sfGFP | This work |
| pSCKtheoRAJ11-oIsa | pSTC2, theoHHAzRAJ11, cisRR12-sfGFP | This work |
| pSCKtheoRAJ12-o11 | pSTC1, theoHHAzRAJ12, cisRAJ11-sfGFP | This work |
| pSCKtheoRAJ12-oIsa | pSTC2, theoHHAzRAJ12, cisRR12-sfGFP | This work |
| pSCKtheoRR12-o11 | pSTC2, theoHHAzRR12, cisRAJ11-sfGFP | This work |
| pSCKtheoRR12-o12 | pSTC2, theoHHAzRR12, cisRAJ12-sfGFP | This work |
| pUAtheoRAJ12 | pIDTSMART, theoHHAzRAJ12, cisRAJ12-LacZα | This work |

